# Analysis and Modeling of Early Estradiol-induced GREB1 Single Allele Gene Transcription at the Population Level

**DOI:** 10.1101/2023.08.30.555527

**Authors:** S. Mahmood Ghasemi, Pankaj K. Singh, Hannah L. Johnson, Ayse Koksoy, Michael A. Mancini, Fabio Stossi, Robert Azencott

## Abstract

Single molecule fluorescence in situ hybridization (smFISH) can be used to visualize transcriptional activation at the single allele level. We and others have applied this approach to better understand the mechanisms of activation by steroid nuclear receptors. However, there is limited understanding of the interconnection between the activation of target gene alleles inside the same nucleus and within large cell populations.

Using the GREB1 gene as an early estrogen receptor (ER) response target, we applied smFISH to track E2-activated GREB1 allelic transcription over early time points to evaluate potential dependencies between alleles within the same nucleus. We compared two types of experiments where we altered the initial status of GREB1 basal transcription by treating cells with and without the elongation inhibitor flavopiridol (FV).

E2 stimulation changed the frequencies of active GREB1 alleles in the cell population independently of FV pre-treatment. In FV treated cells, the response time to hormone was delayed, albeit still reaching at 90 minutes the same levels as in cells not treated by FV. We show that the joint frequencies of GREB1 activated alleles observed at the cell population level imply significant dependency between pairs of alleles within the same nucleus. We identify probabilistic models of joint alleles activations by applying a principle of maximum entropy. For pairs of alleles, we have then quantified statistical dependency by computing their mutual information. We have then introduced a stochastic model compatible with allelic statistical dependencies, and we have fitted this model to our data by intensive simulations. This provided estimates of the average lifetime for degradation of GREB1 introns and of the mean time between two successive transcription rounds. Our approach informs on how to extract information on single allele regulation by ER from within a large population of cells, and should be applicable to many other genes.

**AUTHOR SUMMARY:** After application of a gene transcription stimulus, in this case the hormone 17*β*–estradiol, on large populations of cells over a short time period, we focused on quantifying and modeling the frequencies of GREB1 single allele activations. We have established an experimental and computational pipeline to analyze large numbers of high resolution smFISH images to detect and monitor active GREB1 alleles, that can be translatable to any target gene of interest. A key result is that, at the population level, activation of individual GREB1 alleles within the same nucleus do exhibit *statistically significant dependencies* which we quantify by the *mutual information* between activation states of pairs of alleles. After noticing that frequencies of joint alleles activations observed over our large cell populations evolve smoothly in time, we have defined a *population level stochastic model* which we fit to the observed time course of GREB1 activation frequencies. This provided coherent estimates of the mean time between rounds of GREB1 transcription and the mean lifetime of nascent mRNAs. Our algorithmic approach and experimental methods are applicable to many other genes.

## INTRODUCTION

Time-dependent modulation of gene transcription is necessary for a cell to respond to stimuli in a dynamic and reversible manner. The difficulty in dissecting the complex mechanisms of biological responses is enhanced by the fact that gene transcription in individual cells within a population appears to be vastly heterogeneous (1–8).

We and others have used single molecule RNA FISH (smFISH) to study the effect of stimuli on gene transcription by population analysis of fixed samples (3,7), which facilitates the capture of a large number of events, their spatial location, and the nascent RNA from individual alleles; however, this is at the expense of time dynamics. Specifically, we have focused on the estrogen receptor (ER), a well-established model for transcriptional response to hormones (*i*.*e*., 17*β*-estradiol, E2), using GREB1 as a prototypical early ER target gene (2,8,9). In our previous study (8), we identified that GREB1 responded to E2 in a cell- and allele-dependent manner, and that the frequency of allele activation was tunable by specific epigenetic inhibitors, indicating that the cell has mechanisms in place to control the frequency of allelic responses to stimuli.

To complement the population analysis, several time dynamic studies have been performed. The method of choice has been engineering one or more copies of a gene to contain repeat sequences that are recognized by specific fluorescently-tagged proteins (*e*.*g*., MS2, PP7) (10–12). These live studies proposed that gene transcription occurs in stochastic bursts, where a gene is ON for a short period of time, followed by longer periods of inactivity. Live cell experiments of engineered GREB1 alleles (2), and another prototypical E2-target, TFF1 (1), have shown a highly-dynamic response to hormone in individual cells. From these pivotal studies, the following observations have been made: 1) E2 regulates the frequency of bursting by reducing the promoter OFF times; 2) extrinsic noise governs the cell-by-cell heterogeneity in response; 3) there is some correlation between alleles in the same nucleus; and, 4) at the level of individual cells, live imaging experiments can be accurately fitted by a stochastic model driven by a two-states promoter.

To model the stochastic gene transcription bursts observed in individual cells, several papers (1,2,13-20) have introduced and simulated engineered gene promoter models randomly switching between one active state and several inactive states. The most popular have been two states models where the gene promoter is either ON or OFF. For instance, in Fritzch et al. (2), “two-states” models were fitted to GREB1 transcription bursts observed live in single cells, with model parameters exhibiting quite strong fluctuations from cell-to-cell, which encouraged our present population-level study of endogenous GREB1 gene expression.

Here, we sought to study and model the initial phases of hormone stimulation by using smFISH on fixed MCF-7 breast cancer cells in culture. In our experiments, smFISH images are acquired every 15 minutes, at times *T*_0_ = 0, *T*_1_ = 15 min, …, *T*_6_ = 90 min. At each time *T* = *T*_*j*_, a large cell population pop(T) of N(T) cells is imaged, with N(T) ranging from 400 to 1000. We compared two types of initial conditions:

- in (FV+E2)-experiments, a flavopiridol (FV) pretreatment of cells started two hours before T_0_ to block ongoing transcriptional elongation (21) until FV release at T_0_, which synchronizes the initiation step of the transcriptional cycle.
- in E2-experiments, cells were maintained in a “native” state at T= T_0_, so that the transcription cycle was random before *T*_0_.

At *T*_0_, each experiment started from seven distinct initial cell populations {*init*(0), …, *init*(7)}. Each population *init*(*j*) evolved separately from time *T*_0_ until *T*_*j*_, and the state of *pop*(*T*_*j*_) was only imaged at time *T*_*j*_. This approach does not enable tracking the same cells and active alleles across time. At each time *T* = *T*_*j*_, image analysis of smFISH data provided GREB1 transcription statistics across all cells of *pop*(*T*_*j*_). For *k* = 0,1,2,3,4, we computed the frequencies *Q*_*k*_(*T*) of nuclei exhibiting “k” detectable nascent mRNAs (“active alleles”). The frequencies *Q*_*k*_(*T*) aggregate transcription activities over the *N*(*T*) cells of *pop*(*T*). As seen in (1,2,13-20), transcription bursts of these *N*(*T*) individual cells are *random* and clearly *non synchronous*, so that our transcription data which aggregate GREB1 transcriptions of individual cells across a large population *pop*(*T*) provide population-level activation frequencies which evolve smoothly in time with no significant bursts. To model the smooth time dynamics of the frequencies *Q*_*k*_(*T*), we introduced a “*population-level”* stochastic transcription model involving four key parameters: 1) mean waiting time (“*A*”) between successive productive transcription rounds; 2) mean lifetime (“*L*”) of nascent mRNA; 3) mean elongation time (“*MTD*”) to complete one mRNA; and, 4) the minimal number (“*VTH*”) of RNA molecules enabling fluorescence detection. Parameters estimation for our population-level model was implemented by massive stochastic simulations to reach a good fit to smFISH imaging data across biological replicate experiments. A key step was to determine that at each time *T* = *T*_*j*_, population-level transcription activation frequencies indicated *statistical dependencies* between pairs of GREB1(*AL*_1_, *AL*_2_) alleles within individual nuclei. At each time *T* = *T*_*j*_, we applied a maximum entropy principle to fit a probabilistic model to the frequencies of joint allele activations observed at the population level. This enabled the quantification of dependencies between pairs of alleles by computing their Mutual Information. While the present study focused on GREB1 gene transcription, we expect that the algorithmic modeling and data analysis techniques that we developed will be applicable for other hormone-induced genes transcription.

## RESULTS

### Time course analysis of early E2-induced GREB1 gene transcription by smFISH before and after treatment by flavopiridol

As we have shown previously (8), E2 activates GREB1 gene transcription in a cell- and allele-dependent manner, as measured by smFISH using spectrally separated exon and intron probe sets. Here, we sought to focus on the initial phase of hormonal stimulation (first 90 minutes) by measuring cell population pop(T) responses at 15 minutes intervals (*T*_0_ = 0, *T*_1_ = 15 min, *T*_2_ = 30 min,… up to *T*_2_ = 90 min) under two types of cell state initial conditions and three independent biological replicates per condition type. The first condition was to consider the initial state of gene transcription as random, *i*.*e*., individual alleles have their own “history” with RNA polymerases II located at random phases during either initiation or elongation, which is represented by cells grown for 48 hours in hormone-depleted media. Cell population transcriptional data after this initial cell state are displayed by RED curves and labeled “E2-curves” in every Figure. The second condition was designed to arrest transcription elongation by using the reversible CDK9 inhibitor, flavopiridol (FV), for 2 hours before E2 treatment at *T*_0_, causing elongation to stop and RNA polymerase II to stall at gene promoters (21). We then released the FV block by three washes, and stimulated GREB1 gene transcription by hormone treatment (E2, 10nM); in the Figures the corresponding population data are displayed by BLUE curves and labeled “FV+E2”.

For each independent biological replicate, the analyzed *pop*(*T*) ranged from approximately 400 to 1000 cells, captured by high resolution (60x/1.42NA) epifluorescence deconvolution microscopy (representative images are in **Figure 1A**). Our temporal resolution has limitations as GREB1 probe visualization requires ∼20 individually labeled fluorescent oligo probes (out of 48) to bind to nascent RNA, which, based on their location on the gene, occurs when the GREB1 is ∼60-70% transcribed; for sequences of oligos refer to (8,22). On average, the speed of RNA Polymerase II in mammalian cells (i.e., ∼2-2.5Kb/min, (21,23,24)), suggests that detection of newly made, partial RNAs could occur only after ∼30 minutes of E2 induction.

**Figure 1.**
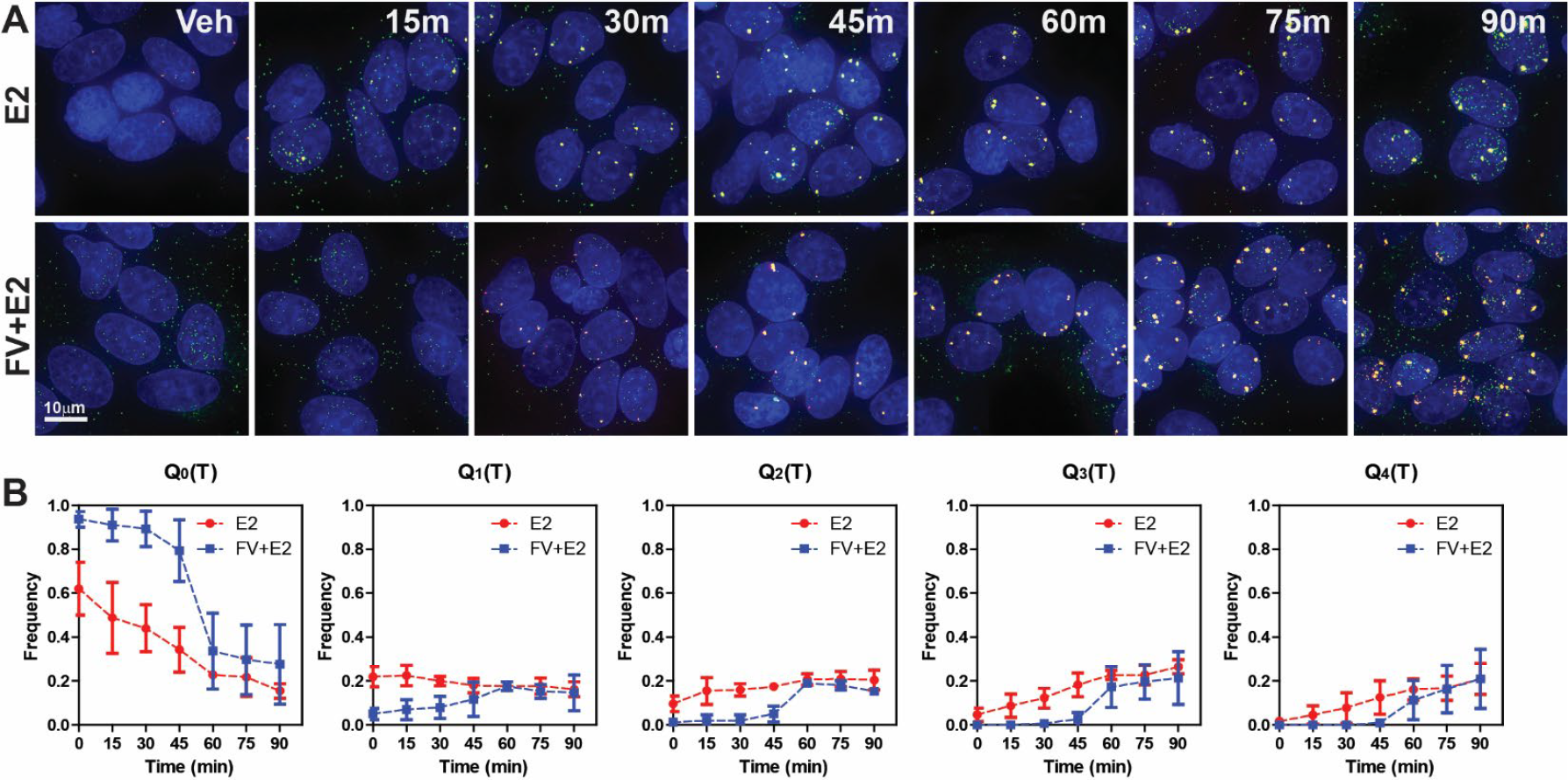
GREB1 smFISH time course analysis in MCF-7 breast cancer cells with and without flavopiridol block/release. A). MCF-7 cells were treated for the indicated times with 10nM E2 and GREB1 smFISH was performed at each time point. Images are at 60x/1.42, deconvolved and max projected. Red spots represent intronic and green spots exonic probe sets. Samples labeled as FV+E2, were pretreated with 1µM flavopiridol (FV) for 2 hours, followed by three washes and E2 treatment. Scale bar: 10µm. B). The time courses of five frequencies Q_0_(T), Q_1_(T), …, Q_4_(T) of active GREB1 alleles/cell are shown as follows. The red curves display the frequencies Q_k_ (T) after averaging over three E2 experiments. The blue curves display the Q_k_ (T) after averaging over three FV+E2 experiments. The vertical bars display the dispersion of Q_k_ values over three similar independent experiments. Note that the dispersion of Q_k_ values across 3 experiments is much larger than the standard error of estimation for Q_k_ (T) in each experiment. At the end of all experiments (T= 90min), all Q_k_ (T) stabilize to a value ≈ 20%.

As smFISH experiments do not directly follow the activation of individual alleles live, the best proxy is to analyze a time series that evaluates transcriptional events across the population *pop*(*T*) of size *N*(*T*) using several statistical approaches describing the set of observable active alleles. In all nuclei, we used custom automated image analysis (described in detail in the Methods section), to identify zero to four active GREB1 alleles [*k* = 0, 1, 2, 3, 4]; from which the number *N*_*k*_(*T*) of nuclei exhibiting k active alleles was calculated, yielding five frequencies, defined by 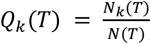. The behavior of the cell population at each time point is then represented by the five GREB1 activation frequencies, *Q*_0_(*T*), *Q*_1_(*T*), *Q*_2_(*T*), *Q*_3_(*T*), *Q*_4_(*T*), which naturally add up to 1 (**Figure 1B**). These frequencies were calculated across more than 400 cells per replicate, with standard error margins of the order of 2.5% (see Supplemental Materials). The effects of flavopiridol block/release on E2-induced gene transcription (**Figure 1B**, blue curves) appear to result in: 1) a significant increase in *Q*_0_(*T*) at all the time points < 60 min; and, 2) a corresponding decrease in the other *Q*_*k*_(*T*), collectively indicating that the FV pre-treatment was effective. More noteworthy are the two following observations: 1) the cell-to-cell and allele-by-allele variation in responses is maintained if transcription elongation is manipulated indicating that this part of the transcription cycle is not controlling synchronicity of responses to hormone; and, 2) the final frequencies of activation at time points > 60 min are virtually identical with or without FV pre-treatment, indicating that the E2 response “catches up” independently of the starting conditions, so that a random cell state starting condition (*i*.*e*., no FV treatment) does not offer an advantage in term of response to hormone over time.

### Allelic activations by E2-induced GREB1 transcription exhibit significant statistical dependency

We explored whether GREB1 activation occurred independently at the four alleles present in each of the aneuploid MCF-7 nuclei by estimating the probabilities of joint activation for pairs of alleles *AL*_1_, *AL*_2_. At time T, in any nucleus *NUC*_***n***_, each allele can either be ON or OFF, therefore yielding 16 distinct possible joint activation states {*S*_0_, *S*_1_, *S*_2_, … *S*_15_} for the 4 alleles *AL*_1_, *AL*_2_, *AL*_3_, *AL*_4_. Denote *prob*_***n***_(*S*_*k*_) as the probability that the four alleles in *NUC*_***n***_ are in the joint state *S*_*k*_. The 16 probabilities *prob*_***n***_(*S*_0_), …, *prob*_***n***_(*S*_15_) add up to 1, and depend on time T. Due to population heterogeneity, *prob*_***n***_(*S*_*k*_) will depend on many cell extrinsic and intrinsic factors specific to each *NUC*_***n***_; hence, our image data could only record averages *F*_*T*_(*S*_*k*_) of the probabilities *prob*_*n*_(*S*_*k*_) over all nuclei *NUC*_***n***_ of pop(T). Concretely, since image resolution did not enable allele matching between distinct cells, the probabilities *F*_*T*_(*S*_*k*_) were not directly computable from image analysis, which could only provide the five observed frequencies *Q*_0_(*T*), …, *Q*_4_(*T*) shown in **Figure 1B**. For each time point T, the algorithmic challenge was hence to compute 16 unknown probabilities *F*_*T*_(*S*_0_), …, *F*_*T*_(*S*_15_) starting only from the 5 observed frequencies *Q*_0_(*T*), …, *Q*_4_(*T*). We identified the five explicit linear relations (see Methods Equation 1) expressing the *Q*_*j*_(*T*) in terms of the *F*_*T*_(*S*_*k*_), but this only provided five explicit linear constraints on our 15 unknown *F*_*T*_(*S*_*k*_). To handle this estimation problem, a natural first approach was to assume that under the probability distribution *F*_*T*_, one had statistical independence of activations among the four GREB1 alleles. In practical terms, independence means that there is no quantifiable mechanism through which any allele interferes or influences activation potential in other alleles in the same nucleus. We have proved (see Methods Equation 2) that, if the probability *F*_*T*_ of joint activations involved statistical independence between alleles activations, the observed frequencies *Q*_0_(*T*), …, *Q*_4_(*T*) would have to verify extremely restrictive polynomial constraints (see Methods MM5 and Equation 2).

All our experiments revealed that these polynomial constraints were never satisfied by the observed *Q*_0_(*T*), …, *Q*_4_(*T*). Thus, to evaluate the *F*_*T*_(*S*_*k*_) from frequencies of joint alleles activations observed at population level, we had to reject the hypothesis of statistical independence between activations of the four distinct alleles within a nucleus.

We have confirmed this theoretical result, by comparing, at each time point, the experimental activation frequencies *Q*_0_(*T*), …, *Q*_4_(*T*) with the analogous activation frequencies generated by joint probability models *F*_*T*_ based on independence between allele activations. Indeed, independence would imply that each probability model *F*_*T*_ is fully determined once one specifies activation frequencies at time T for each single allele *AL*_1_, *AL*_2_, *AL*_3_, *AL*_4_. For each time T, we have generated 10^**6**^ such models F_**T**_ based on the hypothesis of independence between alleles activations, and computed the associated virtual activations frequencies [*virQ*_0_(*T*), …, *virQ*_4_(*T*)] to compare them to the observed *Q*_0_(*T*), …, *Q*_4_(*T*). Our results clearly show that none of these 10^6^ sets of virtual frequencies [*virQ*_0_(*T*), …, *virQ*_4_(*T*)] could match the experimentally observed *Q*_*j*_(*T*). This was visualized by scatter plots, each one displaying a pair [*Q*_*i*_(*T*), *Q*_*j*_(*T*)] observed over time for each experiment. For example, **Figure 2** separately displays scatter plots for the two pairs [*Q*_0_(*T*), *Q*_1_(*T*)] and [*Q*_0_(*T*), *Q*_4_(*T*)]. The red dots represent these pairs of frequencies observed via smFISH or E2 experiments. The blue dots similarly display the same pairs for FV+ E2 experiments. Arrows indicate the successive observations of frequencies [*Q*_0_(*T*), *Q*_1_(*T*)] and [*Q*_0_(*T*), *Q*_4_(*T*)] for an individual experiment. The 10^6^ green dots represent the virtual pairs [*virQ*_0_, *virQ*_1_] or [*virQ*_0_, *virQ*_4_] associated to 10^6^ virtual joint probability models *F*_*T*_ based upon the independence hypothesis. As shown in **Figure 2**, both blue and red dots are positioned well away from green dots, thus confirming that, for population level modeling, one must reject the hypothesis of independence between alleles.

**Figure 2.**
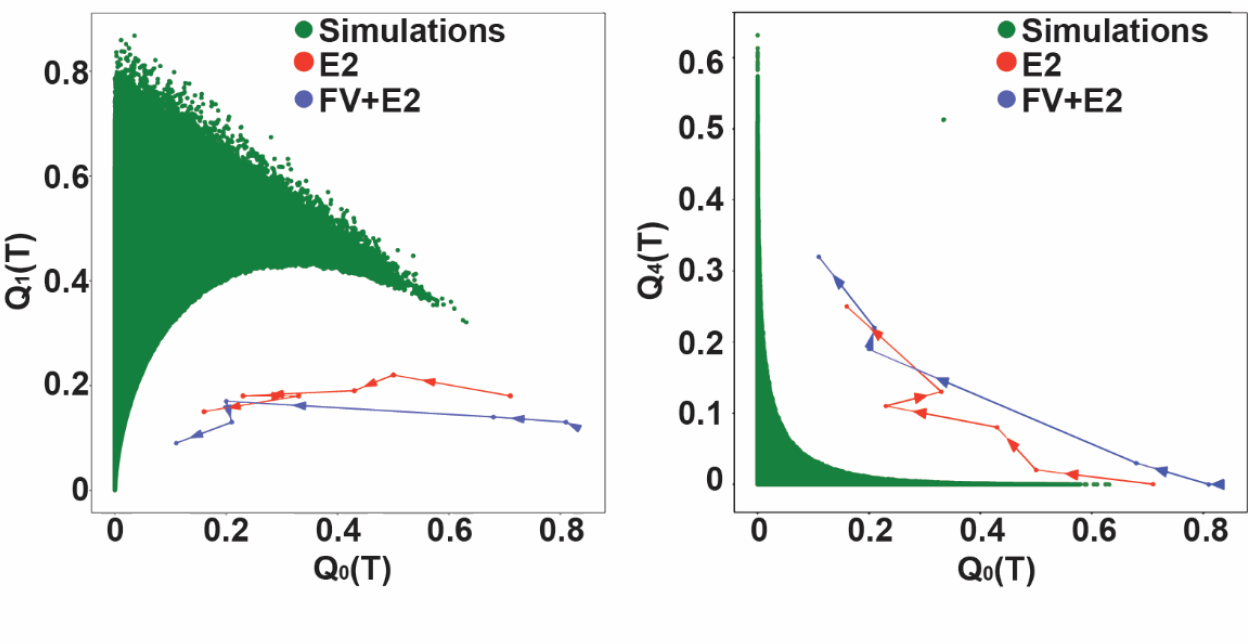
Statistical dependency between GREB1 alleles activations. Scatter plot representation of two frequency pairs (Q_0_(T), Q_1_(T) on the left and Q_0_(T), Q_4_(T) on the right). The blue (FV+E2) and red (E2) curves display the real data time evolutions for the pair of frequencies observed in two experiments. Arrows indicate the time direction. On the left panel, each of the 10^6^ green dots represents one pair of frequencies (q_0_, q_1_) generated by at least one model with independent alleles. Note that the green dots always remain distinct from the experimental red and blue dots. Similar graphs were obtained for any pair Q_i_ (T), Q_j_ (T) with i ≠ j. This indicates that probabilistic modeling of transcription activities aggregated at cell population level requires assuming some dependency between alleles activations.

As just showed above, to be compatible with experimental data, the unknown average frequencies *F*_*T*_(*S*_0_), …, *F*_*T*_(*S*_15_) of jointly activated alleles across cell population *pop*(*T*) must exhibit dependencies between alleles activations. Each one of the 5 observed frequencies *Q*_*j*_(*T*) is an explicit linear combination of the 16 unknowns *F*_*T*_(*S*_0_), …, *F*_*T*_(*S*_15_) (see Methods, Equation 1). Since there was no probabilistic model *F*_*T*_ achieving zero dependencies between alleles activations and also verifying these 5 linear constraints, we decided to seek a model *F*_*T*_ compatible with these 5 constraints and *minimizing dependencies* between alleles activations. For fitting a joint probability distribution *F*_*T*_ to data under linear constraints, a generic principle is that minimizing dependencies is approximately equivalent to *maximizing the entropy Ent*(*F*_*T*_) of the probability model *F*_*T*_ under the same linear constraints (see Methods MM6). This maximum entropy principle is well established in the physics of gases or of spinglass magnets arrays, and has also successfully been used to model images by Gibbs distributions (25,26). Here we have applied this principle to theoretically compute the unique joint probability *F*_*T*_ of activated alleles which has maximum entropy among all probabilities compatible with the five observed frequencies *Q*_0_(*T*), …, *Q*_4_(*T*). We then proved that this max-entropy probability model *F*_*T*_ has full symmetry, meaning that arbitrary permutations of the four alleles do not change the frequencies of their joint state of activations. For instance, due to average probability modeling of the whole population *pop*(*T*), each of the four alleles has the same activation probability 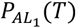 at time T. We have thus obtained explicit formulas expressing each one of the 16 unknowns *F*_*T*_(*S*_0_), …, *F*_*T*_(*S*_15_) in terms of the five observed frequencies *Q*_*j*_(*T*) (see Methods, equation 1). These formulas also gave us explicit expressions for two key probabilities (see Methods, Equation 3), namely: 1) for each single allele *AL*_1_, the probability 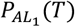 that *AL*_1_ will be active at time T; and 2) for each pair of alleles *AL*_1_, *AL*_2_ in the same nucleus, the probability 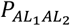 (*T*) that *AL*_1_ and *AL*_2_ will be simultaneously active at time T. The probabilities 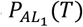 and 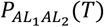 do not change if we replace *AL*_1_, *AL*_2_ by any other two alleles *AL*_*i*_, *AL*_*j*_ within the same nucleus. This is due to the full symmetry of the joint probabilities *F*_*T*_(*S*_*k*_), a property which was derived from the maximum entropy principle.

In **Figure 3 A**, we display the evolution of 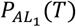 over time. For FV+ E2 experiments (blue curve), the initial 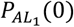 is nearly 0 and 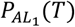 remains practically equal to zero until *T* = 30 min since GREB1 transcription duration is of the order of 40 min and GREB1 transcriptions are nearly blocked by FV before *T* = 0. For E2 experiments (red curves), 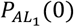 is naturally higher than for FV+E2 experiments (blue curves) due to some GREB1 transcription activity at low level before infusion of E2 at *T* = 0. For all experiments, 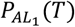 increases steadily with T, and reaches a maximum ranging from 40% to 60% at *T* = 90 min.

**Figure 3.**
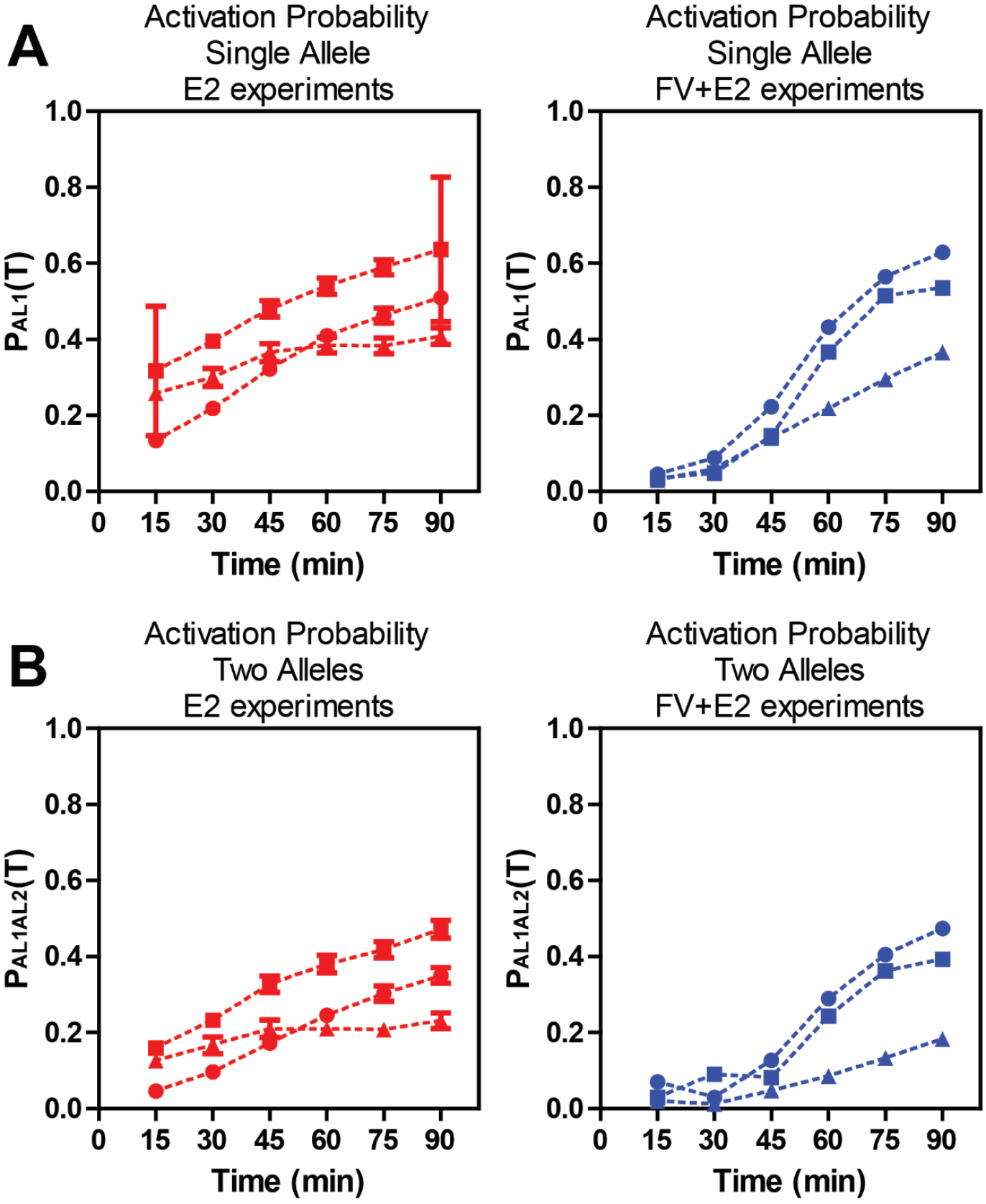
Time course for GREB1 activation probabilities 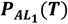 and 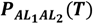. A) Here 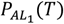 is the computed probability that a given single allele AL_1_ will be GREB1 activated at time T, and B) 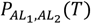 is the joint probability that a given pair of alleles AL_1_, AL_2_ will both be activated at time T. The time courses of 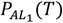 and 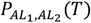 between T = 15 min and T = 90 min are displayed by red curves for three E2 experiments and by blue curves for three (FV+E2) experiments. The vertical bars display the standard errors on these probabilities.

For each pair of alleles (*AL*_1_, *AL*_2_) in the same nucleus, their joint random activation states at time T can have only one of four possible configurations: active/active, active/inactive, inactive/active, or inactive/inactive. We have explicitly computed the probabilities of these four configurations in terms of the observed frequencies *Q*_*j*_(*T*). The probability 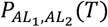 that the two given alleles *AL*_1_ and *AL*_2_ are simultaneously active at time T was then plotted for all experiments (see **Figure 3B**). For FV+ E2 experiments (blue curves), 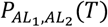 remains quite low until *T* = 30 min since most polymerase elongations started after time *T* = 0 are still too incomplete at *T* = 30 min to be reliably detectable. For all experiments, 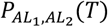 increases with T, and reaches a maximum ranging from 25% to 45% at *T* = 90 min.

### Statistical Validation of Dependency between GREB1 alleles activations after E2 treatment

For our probabilistic model *F*_*T*_ fitted by maximum entropy to actual GREB1 activation frequencies aggregated at the cell population level, our explicit computation of the probabilities 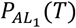 and 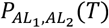 enabled us to test whether the activation of alleles *AL*_1_, *AL*_2_ is statistically dependent of each other, and to quantify their statistical dependency. Indeed at time T, statistical independence for the activation of alleles *AL*_1_, *AL*_2_ would classically imply the equality 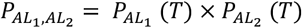. In our experiments this equality is *significantly not satisfied*, as validated by our detailed analysis of estimation errors on 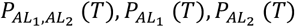 (see Methods MM5,MM6).

At time T, the conditional probability that {*AL*_2_ *is active*} given that {*AL*_1_ *is active*} is classically computed by 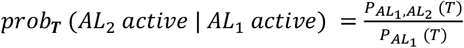.

For all our 6 experiments (see **Figure 4**), we have 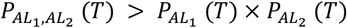 at each time point *T* ≥ 15 min. This forces the conditional probability *prob*_*T*_(*AL*_2_ *active*| *AL*_1_ *active*) to be always larger than the unconditioned probability 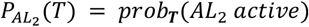. This is a clear indicator of statistical dependency between the activations of *AL*_1_ and *AL*_2_. Indeed one can quantify the level of dependency between activations of *AL*_1_ and *AL*_2_ by comparing the **dependency ratio** 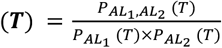. to the baseline value 1. Our detailed statistical study of estimation errors on *dep*(*T*) (see Methods section MM6) shows that *the inequality* {*dep*(*T*) < 1} *is significantly valid* at the 95% confidence level for all our E2 experiments as soon as *T* ≥ 15 min, and for all our FV+E2 experiments when *T* ≥ 30 min. In this time range this proves *significant statistical dependency* between the activations of any alleles pair *AL*_1_, *AL*_2_. In fact, the dependency ratio *dep*(*T*) remains larger than 1.15 during the times analyzed. Hence, when *AL*_1_ is active at time T, the conditional probability that *AL*_2_ is also active is at least 15% higher than the unconditioned activation probability for *AL*_2_. Note that for initial times *T* = 0 or *T* = 15 min, the probabilities 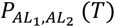 and 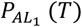 are typically too small for reliable statistical estimation of the ratio *dep*(*T*).

**Figure 4.**
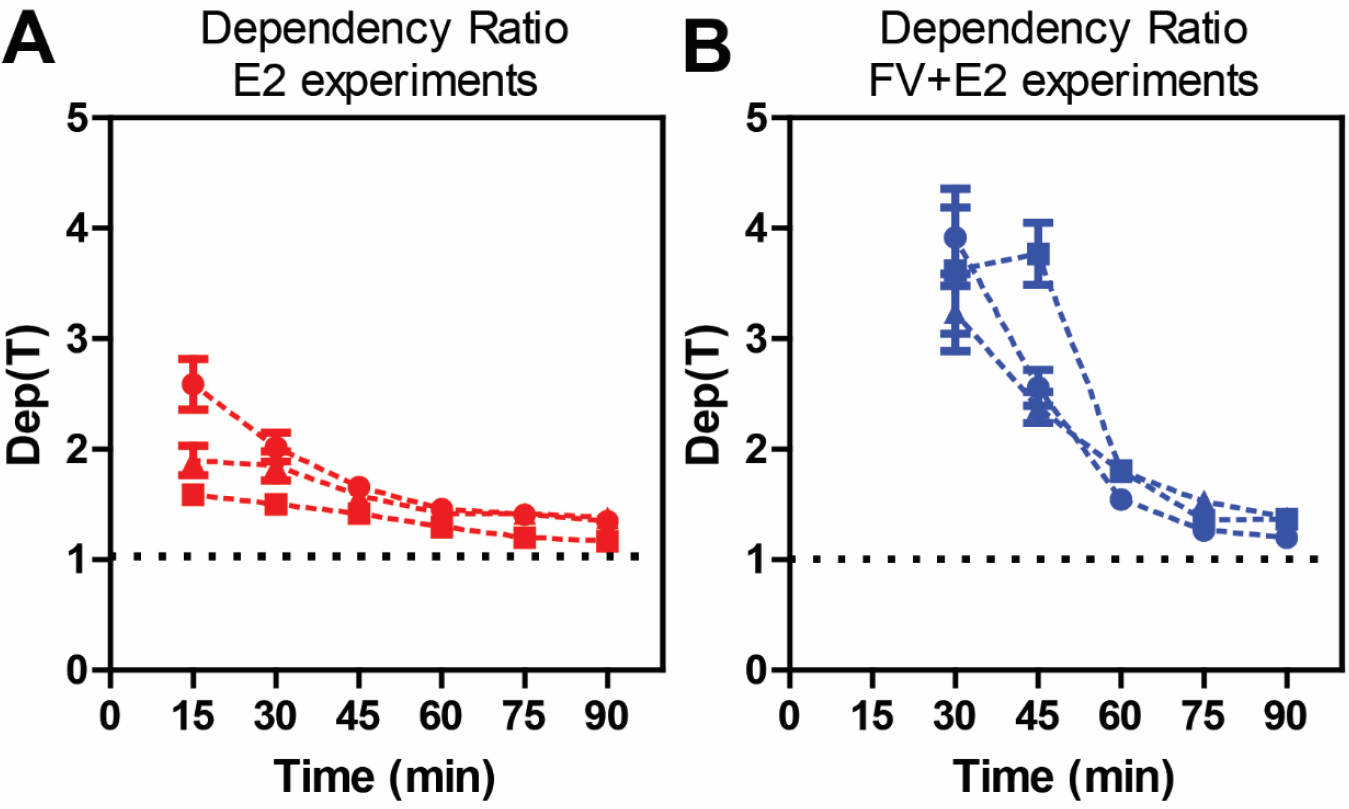
For pairs of alleles AL1, AL2, the dependency ratio 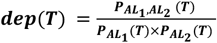 is significantly larger than 1 at confidence level 95%, which indicates significant statistical dependency between GREB1 activations of AL_1_ and AL_2_. We display the time course dep(T) at all T ≥ 15 min for three E2 experiments (A) and at all T ≥ 30 min for three FV+E2 experiments (B), The vertical bars display the standard errors of estimation on dep(T). Note that the ratio dep(T) remains larger than 1.15 in these time ranges.

The probabilistic dependency between the activation states of two alleles *AL*_1_, *AL*_2_ can also be quantified by their *Mutual Information* 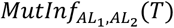 (see formulas in Methods section MM6). Recall that 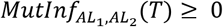 evaluates how knowing that *AL*_1_ is active at time T improves the accuracy of predicting if *AL*_2_ is active at time T. Complete independence of *AL*_1_ and *AL*_2_ would imply 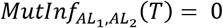, so strictly positive values of 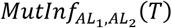 indicate dependency between the activation of *AL*_1_ and *AL*_2_. Since the population average probability *F*_*T*_ of jointly activated alleles has full symmetry, all allele pairs *AL*_1_, *AL*_2_ must have mutual information identical to 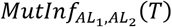. We have computed 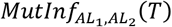 for all experiments, and all T. As detailed in Methods section MM6, the generic formula giving 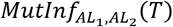 involves terms such as 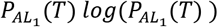 and 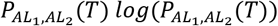 for which the estimation errors become high when the activation probabilities 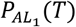 and 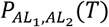 are very small. For (FV+E2) experiments, and for *T* ≤ 30 min, both 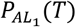 and 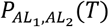 are very close to 0, so that the natural estimates of 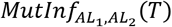 become statistically reliable only for *T* ≥ 45 min. For E2 experiments, since GREB1 transcription activity starts before *T* = 0, one can reliably estimate 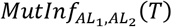 as soon as *T* ≥ 15 min. In **Figure 5**, we display the time course of mutual information 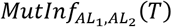 for each one of our experiments.

**Figure 5.**
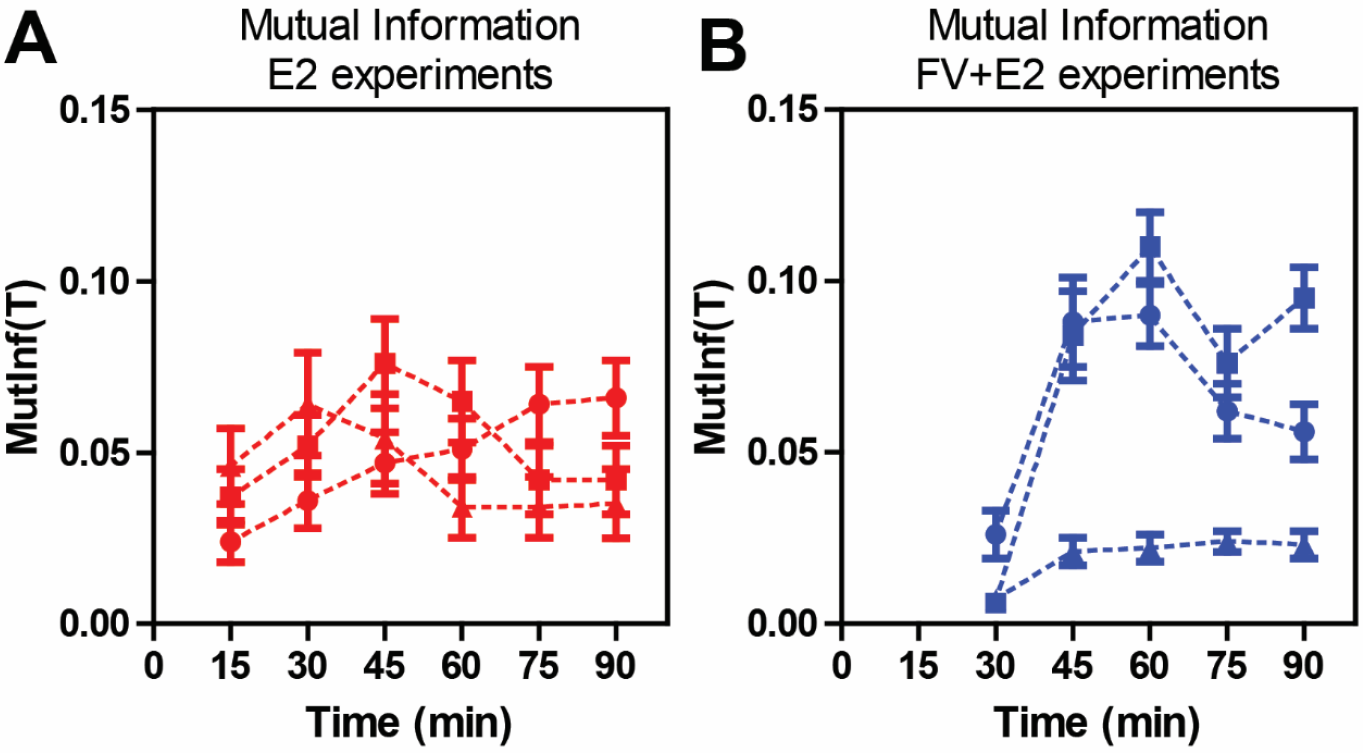
Mutual Information between pairs of alleles. For each one of our experiments, and at all time points T, we have computed the Mutual Information 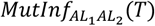 between the stochastic GREB1 activations of any given pair of alleles AL_1_, AL_2_ within the same nucleus. We display the time course of 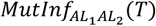 for three E2 experiments and T ≥ 15 min (A) as well as for three FV+E2 experiments and T ≥ 30 min (B). The vertical bars display the standard errors on 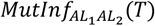, and show that in these time ranges, the mutual information is always significantly positive with confidence level 95%, which indicates a statistically significant level of dependency between GREB1 activations for pairs of alleles AL_1_, AL_2_. At times T = 0, T = 15 min for FV+E2 experiments, and at T = 0 for E2 experiments, the joint probabilities 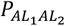 are too small to accurately compute 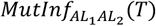.

For FV+E2 experiments as well as for E2 experiments, and for 45 *min* ≤ *T* ≤ 90 *min*, 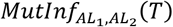 roughly ranges from 0.04 to 0.09. For our ranges of mutual information values 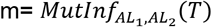, the dependency ratio *dep*(*T*) can be roughly approximated by 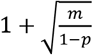, where 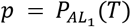. This mathematical approximation valid for small “m” explains why small mutual information values reflect much more sizeable positive values for the difference (*dep*(*T*) − 1). The estimation errors indicate that for 45 *min* ≤ *T* ≤ 90 *min* and for all our experiments, the mutual information 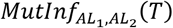 is *significantly positive* at the 95% confidence level. This confirms that, at the population level, we detect significant statistical dependency between jointly activated allele pairs within the same nucleus.

### Population level stochastic model to emulate time course of GREB1 allele activation frequencies

We next sought to fit a *population level* stochastic model dedicated to emulating the dynamics of GREB1 transcriptional frequencies computed across large cell populations. Several papers (see 2,18,20) have modeled the dynamics of random gene transcription bursts observed in live *single cells* by stochastic “two-states” promoter models, in which gene promoters are viewed as stochastic automatons randomly cycling through an ON-state and an OFF-state. In these studies, the parameters of two-states promoter models are separately fitted to each single cell continuously observed at very short time intervals. As explicitly pointed out by (2), the estimated parameters of these single cell models vary quite strongly (up to 20%) from cell to cell, due to heterogeneities in cells biology and/or in their local chemical environment. In our smFISH experiments, the frequency *Q*_*k*_(*T*) of nuclei exhibiting “*k*” GREB1 activated alleles at time T is estimated by averaging across several hundred cells of *pop*(*T*). Since random gene transcriptions bursts are *highly decorrelated* from cell to cell and have short duration, averaging joint activations frequencies across *pop*(*T*) essentially smooths out the random GREB1 transcription bursts occurring in single cells. We have verified this intuitive point by simultaneous simulation of *N* = 400 “two-state” promoter models for GREB1 transcription, followed by averaging at each time T the GREB1 transcription bursts occurring at time T among these N simulations. In our experiments, which involve large cell populations, the observed frequencies *Q*_*k*_(*T*) indeed have rather smooth time evolutions, as well as the probabilities 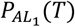 and 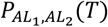 derived from the frequencies *Q*_0_(*T*), …, *Q*_4_(*T*).

To emulate the time course of GREB1 transcription frequencies observed across each cell population *pop*(*T*), we introduce a *population level* stochastic model, where successive ER DNA binding occurs randomly after exponentially distributed waiting times that can be followed by coregulator recruitment and transcription initiation. At these random transcription initiation times, GREB1 mRNA elongation proceeds with fixed *Mean Transcription Duration (MTD)*. Several studies (21,23,24) indicate that gene transcription occurs at a roughly constant speed of ∼2 − 2.5 *krob*/*min*, which results in an *MTM* ≃ 44min for GREB1. Random time durations between successive rounds of GREB1 mRNA elongation are assumed to be independent of each other, and to have the same exponential density with *mean value A*, which is a model parameter. Such random time gaps are characteristic of Poisson stochastic processes.

We denote “*nas*” any complete nascent GREB1 mRNA, and “*exonas*”/“*intnas*” the *exonic* and intronic parts of *nas*. The (random) lifetimes of *exonas* and intnas are assumed to have exponential decay. The mean half-life of *exonas* has been empirically calculated via actinomycin D pulse-chase experiments and is approximately 3 hours. As our experiments last 90 minutes, the *exonas* decay does not significantly affect *nas* visibility during this time. However, the intronic component *intnas* was calculated to have a mean lifetime < 35 min, which does directly affect the lifetimes of completed nascent mRNAs. The random lifetime of any complete nascent GREB1 mRNA (from completion to nearly full decay) is hence assumed to have an exponential density with unknown *mean value L*.

Our population level model is thus determined by 3 unknown parameters {*A, L, MTM*}. Since analysis of our smFISH images suggest that the smallest nascent mRNA spots may not be reliably detected, we introduce another unknown parameter, the *Visibility Threshold (VTV)* such that nascent mRNA spots are detectable on our images only if they contain at least *VTV* molecules. For any plausible values of {*A, L, MTD, VTH*}, this model enables rapid simulations generating frequencies 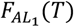 of a single allele activation at time T. Quality of fit is evaluated by the differences 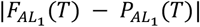 over a range of time points T, where the probability 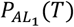 is computed as above from the frequencies *Q*_0_(*T*), …, *Q*_4_(*T*) computed via analysis of smFISH images.

We point out that our population level stochastic model does not attempt to model GREB1 transcriptions in *single cells*. Indeed, our model aims only to emulate the allele activation frequencies resulting from the *aggregation (at cell population level)* of the random allele activations generated by several hundreds of independent two-states stochastic models of GREB1 transcription activities, with two-states model parameters varying slightly from cell to cell.

### Estimation of parameters for our population level model by intensive simulations

To fit the population level model to GREB1 transcription data provided by each FV+E2 experiment, we had to estimate the four parameters {*A, L, MTD, VTH*} which were expected to belong to naturally pre-defined ranges (see Methods, section MM8). But for E2 experiments with no flavopiridol pretreatment, spontaneous GREB1 transcription events at low rates can start long before E2 treatment at *T* = 0. Some of the alleles activated within the last hour before *T* = 0 will generate incomplete nascent mRNAs that will not be detectable at T = 0 and will only be detected by image analysis after *T* = 15 min or *T* = 30 min. Taking into account these nascent mRNAs whose transcription started before *T* = 0 complicates the data analysis for E2 experiments with no FV pretreatment, and requires introducing a new parameter, namely the mean value *A*^+^ of waiting times between successive GREB1 transcription rounds before E2 treatment. As shown in (1,2), E2 treatment increases the frequency of genes transcriptions, so we should assume that *A*^+^ ≥ *A*. Thus, fitting population level data for E2 experiments with no FV pretreatment requires the estimation of five parameters {*A*^+^, *A, L, MTD, VTH*}.

For the unknown values of the parameters {*A*^+^, *A, L, MTD*}, natural ranges were identified from existing literature (see Methods, section MM8, MM9). For the small integer *VTH*, the potential range of values was evaluated by analyzing the rough number of molecules within mRNA smFISH spots detected in the images.

To actually fit the parameters of our population level models to each GREB1 experiment, we performed intensive simulations of this stochastic model to systematically explore the full discretized ranges of the 5 parameters {*A*^+^, *A, L, MTD, VTH*}. For each combination of parameters values, and each time point T, these simulations yielded estimates for the activation frequency 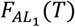 of single alleles across a large population of virtual cells. We only retained the model parameters with good fit to data, *i*.*e*., ensuring that 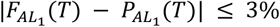, where the probability 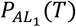 of single allele activation was derived as above from image analysis. We selected final parameters values by enforcing parameter stability across all 6 experiments for *L, MTD, VTH*. Tables 2 and 3 display the best fit parameter values for three experiments with no FV pre-treatment, and for three FV pre-treated experiments. Our best model parameters achieved a quality of fit ≈ 3% for each experiment, in good compatibility with the margins of error on the 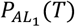 derived from image data.

**Table 1.**
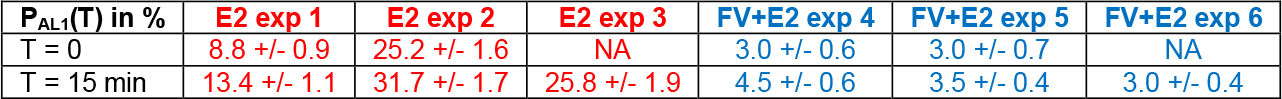
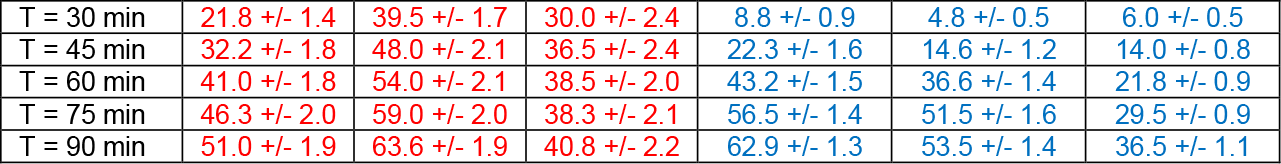
Activation probability 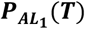 **for single allele *AL***_1_.

**Table 1b.** Joint activation probability 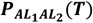 for two alleles *AL*_1_, *AL*_2_.

| $P_{AL_1 AL_2}(T)$ in % | E2 exp 1 | E2 exp 2 | E2 exp 3 | FV+E2 exp 4 | FV+E2 exp 5 | FV+E2 exp 6 |
| --- | --- | --- | --- | --- | --- | --- |
| T= 0 | 2.8 +/- 0.6 | 11.4 +/- 1.3 | NA | 0.5 +/- 0.2 | 0.3 +/- 0.2 | NA |
| T= 15 min | 4.6 +/- 0.7 | 16.0 +/- 1.5 | 12.6 +/- 1.6 | 0.7 +/- 0.2 | 0.3 +/- 0.1 | 0.2 +/- 0.1 |
| T= 30 min | 9.6 +/- 1.1 | 23.3 +/- 1.7 | 16.7 +/- 2.2 | 3.0 +/- 0.6 | 0.9 +/- 0.2 | 1.2 +/- 0.2 |
| T= 45 min | 17.2 +/- 1.6 | 32.7 +/- 2.2 | 21.0 +/- 2.3 | 12.7 +/- 1.4 | 8.1 +/- 1.0 | 4.7 +/- 0.5 |
| T= 60 min | 24.5 +/- 1.7 | 38.0 +/- 2.3 | 21.0 +/- 1.9 | 29.0 +/- 1.4 | 24.3 +/- 1.4 | 8.5 +/- 0.7 |
| T= 75 min | 30.2 +/- 2.0 | 41.8 +/- 2.2 | 20.8 +/- 1.9 | 40.5 +/- 1.5 | 36.2 +/- 1.7 | 13.3 +/- 0.7 |
| T= 90 min | 35.0 +/- 2.0 | 47.2 +/- 2.3 | 23.1 +/- 2.1 | 47.4 +/- 1.5 | 39.3 +/- 1.4 | 18.3 +/- 1.0 |

**Table 2.**
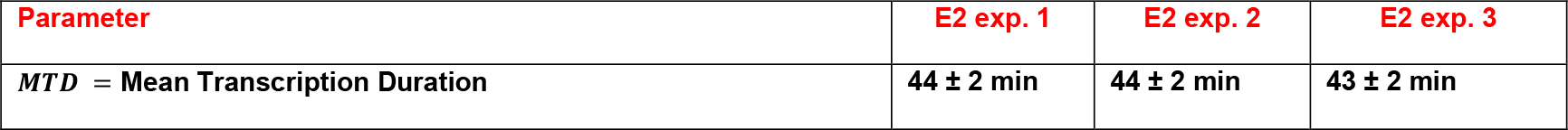
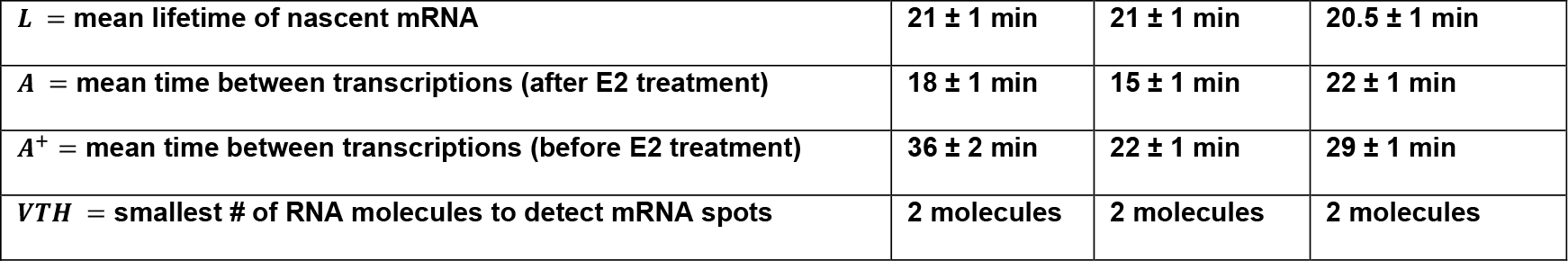
E2 experiments: Parameters estimates for three Population Level Models:

**Table 3.** FV+E2 experiments: Parameters estimates for three Population Level Models.

| Parameter | FV+E2 exp. 4 | FV+E2 exp. 5 | FV+E2 exp. 6 |
| --- | --- | --- | --- |
| $MTD$ = Mean Transcription Duration | $44 \pm 1$ min | $44 \pm 1$ min | $44 \pm 2$ min |
| $L$ = mean lifetime of nascent mRNA | $21 \pm 1$ min | $20.5 \pm 1$ min | $20.5 \pm 1$ min |
| $A$ = mean time between transcriptions (after E2 treatment) | $14.5 \pm 1$ min | $16 \pm 1$ min | $23.5 \pm 1$ min |
| $VTH$ = smallest # mRNA molecules to detect mRNA spots | 2 molecules | 2 molecules | 2 molecules |

These two tables exhibit good stability across all 6 experiments for the mean transcription duration (*MTD* ≃ 44 *min*), the mean lifetime (*L* ≃ 21 *min*) of nascent mRNAs, and the number (*VTH* = 2) molecules of mRNA necessary to detect a nascent mRNA spot. The mean waiting time A between successive GREB1 transcription cycles after E2 treatment had a wider range between 15 min and 23 min among all 6 experiments. We have not identified the main factors influencing the waiting times A, but cell population heterogeneity is likely to strongly impact the variations in A observed from one experiment to the next. Indeed in (2), the authors mentioned that for their two-states model focused on GREB1 transcription observed on separate single cells, the estimated model parameters (such as our parameter A) did strongly vary from cell to cell, with relative variations up to 20%. It is also possible that A may not remain strictly constant in time during E2 induction.

## DISCUSSION

Stimulus-controlled gene transcription is one of the essential ways a cell senses and responds to environmental changes. While this process has been heavily studied in multiple models, a full understanding of how the regulation of events leading to gene transcription unfolds is constantly evolving. From numerous studies across species and models (1–8,11,14,16,18,19), it appears that cells respond to stimuli in a very heterogeneous manner and, even within the same nucleus, different copies of the same target gene respond asynchronously. Is this because of fully stochastic biological reactions, or given the evolutionary development of regulated mechanistic steps in gene expression, is regulated allelic activation a way to finely-tune individual cell responses to external cues? In our earlier study (8), we suggested cells can use an epigenetic mechanism to control the frequency of active alleles in the nucleus. Here, we focused on the same biological system, the hormone (E2) stimulated GREB1 gene (2,8,9) in MCF-7 breast cancer cells to ask a few additional basic questions: 1) can we synchronize the response of individual alleles by altering transcription elongation? 2) can we determine if alleles in the same nucleus are acting independently or not? 3) can we develop a simplified model to emulate, at the cell population level, the first phases of hormonal response over time, with stability of the model parameters across independent biological replicates?

To address these questions, we compared GREB1 transcription activity in large cell populations under two types of initial conditions: 1) FV +E2 experiments where prior to E2 treatment of our cell populations at *T* = 0, transcription elongation was synchronized and then restarted by addition and wash-out of the reversible CDK9 inhibitor, flavopiridol (FV). 2) E2 experiments where at *T* = 0, cell populations are still in their natural random state after several hours without hormone treatment and with their transcription cycles left untouched. Our experimental data strongly indicate that “synchronizing” RNA Polymerase II at the elongation step is not sufficient to synchronize hormonal responses at the cell-by-cell or allele-by-allele levels. Indeed, for the two types of initial cell population conditions, at the end time point (90 minutes post E2), identical values are reached by key characteristics such as the activation probability of each single allele and the joint activation frequencies for pairs or triplets of alleles.

At each time T, our detailed analysis of observed frequencies for alleles jointly activated by GREB1 nascent mRNA spots demonstrated a *significant statistical dependency* between pairs of activated alleles within the same nuclei. This led us to apply, at each time T, a principle of *maximum entropy under constraints* to compute a probability distribution *F*_*T*_ for the joint activation states of the four alleles within typical nuclei. We then used the joint probability *F*_*T*_ to compute the mutual information 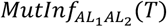 between GREB1 activations of alleles *AL*_1_ and *AL*_2_, in order to quantify the dependency between pairs of alleles. A detailed error analysis for the estimated 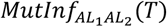 showed that this mutual information had statistically significant positivity for all *T* ≥ 30 min, a clear indicator of moderate but significant dependency between activations for pairs of alleles. An interesting still open question is to identify biochemical factors enabling these dependencies, such as extrinsic chemical factors that can jointly affect all 4 alleles in each nucleus. Other cell-linked factors affecting GREB1 transcription of all 4 alleles were also invoked in (2) to explain the high variation of transcription model parameters fitted separately to single cells.

The probability distributions *F*_*T*_ were computed at each fixed time T from population level frequencies of joint alleles activations. To emulate the time dynamics of these probabilities *F*_*T*_ across time, we introduced a “population level” stochastic model, where random initializations of GREB1 transcriptions are driven by a Poisson process, and are always followed by actual elongation. We were led to introduce this *population level* model instead of the popular two-states models used for *single cell* transcription data (1,2) because averaging GREB1 transcription activity across large cell populations strongly smooths out the random transcription bursts occurring independently among individual cells. Since our smFISH image acquisition modalities do not enable the monitoring across time of single cells GREB1 transcription activity, we designed our population level model to roughly emulate the superposition of several hundreds of independent two-states models of single cells gene transcription dynamics. For each one of our experiments (three FV+E2 experiments and three E2 only experiments) the parameters of our population level model were fitted to experimental data by intensive simulations exploring a very large set of combined parameter values. After this fitting of model parameters to data, the quality of fit was quite precise (less than 3% error on emulated frequencies of GREB1 activations), and across all 6 experiments we achieved good stability for the main parameters, namely the mean elongation duration (*MTD* ≃ 44 *min*), the mean lifetime (*L* ≃ 21*min*) of nascent mRNAs, and the number (*VTH* = 2) of mRNA molecules necessary for reliable detection of a nascent mRNA spot. The estimated mean waiting time A between successive GREB1 transcription after E2 treatment had a wider range (15min to 23min) among our 6 experiments. This variation is quite compatible with the 20% range of variation reported in (2) for the parameters of two-states models fitted separately to single cells.

We expect our innovative modeling approach for hormone-regulated target gene activity observed at population level to be applicable for many other genes and stimuli, a point we intend to validate through further experiments. An interesting and open challenge is to concretely identify the main cell-dependent factors simultaneously impacting transcriptional responses at individual alleles within the cell nucleus.

## MATERIALS AND METHODS

### MM1. Cell culture, materials and treatments

MCF-7 cells were obtained from BCM Cell Culture Core, which routinely validates their identity by genotyping; cultures are constantly tested for mycoplasma contamination as determined by DAPI staining. MCF-7 were maintained in MEM plus 10%FBS media, as recommended by ATCC, except phenol red free and kept in culture for less than 60 days before thawing a fresh vial. Three days prior to experiments, cells were plated on round poly-L-lysine coated coverslips in media containing 5% charcoal-dextran stripped and dialyzed FBS-containing media. Treatments with 17β-estradiol (E2, Sigma) were performed as in (8). For flavopiridol (FV+E2) experiments, cells were treated with FV 1µM for 2 hours, then removed and cells were washed 3x with media prior to E2 1nM treatment for the indicated times.

### MM2. Single molecule RNA FISH (smFISH)

GREB1 smFISH was performed as described in detailed protocols (8,22). Briefly, cells were fixed with 4% paraformaldehyde in PBS, on ice for 20 min. After a PBS wash, cells were left in 70% ethanol for a minimum of 4 hours prior to hybridization (o/n, 37C) with the previously validated GREB1 probe sets (LGC Biosearch Technologies) covering introns (Atto647N) and exons (Quasar570) of the GREB1 gene.

### MM3. Imaging

High resolution imaging for smFISH was performed on a Cytivia DVLive epifluorescence image restoration microscope with an Olympus PlanApo 60×*/*1.42NA objective and a 1.9k × 1.9k sCMOS camera. Z stacks (0.25 µm) covering the whole nucleus (∼10 µm) were acquired before applying a conservative restorative algorithm for quantitative image deconvolution. Ten or more random fields of view (FOVs) were acquired for each time point.

### MM4. Image analysis

Each FOV has three fluorescence channels (DAPI, Q570 (exons) and A647N (introns)) in a 3D-image of size ≃ 1780 × 1780 × 25. Each 3D image channel was projected on its maximum intensity horizontal layer and then analyzed as a 2D image. In the DAPI channel, we first detect and identify cell nuclei. The main steps are: contrast thresholding, connected components detection, elimination of holes, and size filtering. After dilating the detected nuclear mask, we estimate cytoplasm boundaries by the “watershed” segmentation algorithm.

Classical image segmentation techniques are applied in the two other channels to separately detect exonic and intronic spots. Contrast analysis is implemented by OTSU thresholding (27) for the exonic channel (Q570), and by max-entropy thresholding (28) for the intronic channel (A647N). We tested moderate variations of these thresholds to define accuracy margins for these two key spots detections. Nascent mRNAs spots are detected within each nucleus by identifying all pairs (exonic spot + intronic spot) having non-empty intersection. At each time point, the number of active alleles detected per nucleus ranges from 0 to 4 since each cell has 4 GREB1 alleles (29). Segmentation errors due to local cell packing or nuclei overlaps may occasionally generate spurious detections of more than 4 activation spots in a very small percentage of nuclei, which are then automatically discarded. Mature RNAs are identified as exonic spots (Q570) located within the cytoplasmic mask.

### MM5. Probability F_T_ of joint alleles activations at fixed time T

A key modeling step was, at each fixed time T, to estimate the probabilities of joint alleles activation for pairs, triplets, quadruplets of alleles within a cell. We could only aim to estimate averages over *pop*(*T*) for each one of these joint probabilities, since in our experimental setup, distinct alleles are not identifiable. At time T, in any given cell, each allele *AL*_*j*_ (*j* = 1,2,3,4) can either be active (ON state “1”), or not (OFF state “0”). The joint state of the four alleles [*AL*_1_, *AL*_2_, *AL*_3_, *AL*_4_] is then described by a four-digit binary code, with 2^**4**^ = 16 possible joint states labeled

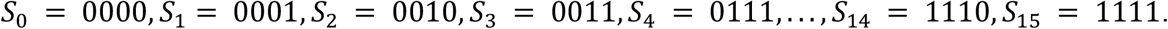

At time T, the cell population *pop*(*T*) contains *N* = *N*(*T*) nuclei, denoted *NUC*_**1**_, *NUC*_**2**_, …, *NUC*_***N***_. For each nucleus *NUC*_***n***_, the current joint state for the four alleles will be equal to *S*_*k*_, with some unknown probability *prob*_***n***_(*S*_*k*_). The 16 probabilities *prob*_***n***_(*S*_0_), …, *prob*_***n***_(*S*_15_) add up to 1, and also depend on the time point T. But due to biochemical heterogeneity of the cells in the population at time T, the *prob*_***n***_(*S*_*k*_) may also depend on unknown biochemical factors specific to nucleus *NUC*_***n***_. Since our frequencies *Q*_*k*_(*T*) of nuclei exhibiting *k* activated alleles at time T are computed across the whole of *pop*(*T*), they only provide reliable information on the average *F*_*T*_(*S*_*k*_) of the probabilities *prob*_***n***_(*S*_*k*_) over all nuclei indices *i* = 1 … *N*. More precisely, for each possible joint state *S*_*k*_ of the four alleles, we want to estimate the average probability 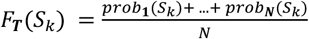.

In a perfectly homogeneous cell population *pop*(*T*), the probabilities *prob*_***n***_(*S*_*k*_) would not depend on the nucleus *NUC*_***n***_ at all, and would hence also be equal to the average probability *F*_*T*_(*S*_*k*_). This ideal situation is blurred by the significant biological diversity of GREB1 transcriptional response from cell to cell, a point well documented in (2).

The frequencies*F*_*T*_(*S*_*k*_) are not directly observable, since smFISH images do not enable specific matching of alleles *AL*_1_, *AL*_2_, *AL*_3_, *AL*_4_ from cell to cell. But as mathematically detailed in Methods and in Suppl. Materials 1 the observed activation frequencies *Q*_*k*_(*T*) are linked to the unknown frequencies

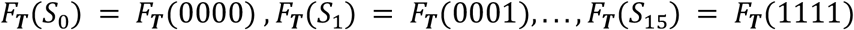

by the following five linear relations.

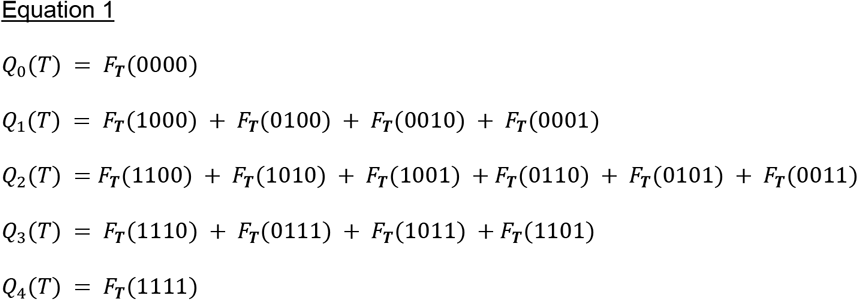

Since these five linear relations cannot determine the 16 unknown probabilities *F*_*T*_(*S*_*k*_), we first checked if we could assume independence of alleles activation. Under the probability F_**T**_ of joint alleles activations, denote *M*_***j***_(*T*) the probability that the specific allele *AL*_*j*_ is activated. Assume temporarily that under *F*_*T*_ the activations of each allele *AL*_1_, *AL*_2_, *AL*_3_, *AL*_4_ are statistically independent of each other (*i*.*e*., there are no mechanisms through which alleles interfere / influence each other’s activations). Independence implies that each unknown joint probability *F*_*T*_(*S*_*k*_) can be expressed directly in terms of *M*_**1**_(*T*), *M*_**2**_(*T*), *M*_**3**_(*T*), *M*_**4**_(*T*) by simple product formulas such as *F*_*T*_(0101) = (1 − *M*_**1**_(*T*)) *M*_**2**_(*T*) (1 − *M*_**3**_(*T*)) *M*_**4**_(*T*), *F*_*T*_(1110) = *M*_**1**_(*T*) *M*_**2**_(*T*) *M*_**3**_(*T*) (1 − *M*_**4**_(*T*)), …, etc. Combining these product formulas with equation 1 we have formally proved (see Suppl. Materials 2) that statistical independence of single allele activations under the joint probability F_**T**_ forces the following polynomial equation of degree 4

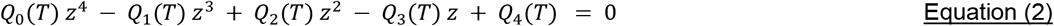

to have four positive and real valued roots *z*_1_, *z*_2_, *z*_3_, *z*_4_. We also showed that

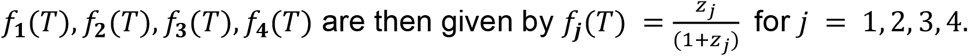

Requiring a polynomial of degree 4 to have four positive and real valued roots *z*_1_, *z*_2_, *z*_3_, *z*_4_ imposes very restrictive constraints on the polynomial coefficients *Q*_0_(*T*), …, *Q*_4_(*T*). In Suppl. Materials 2, we give examples of these polynomial constraints. Our experiments showed very clearly that these constraints are never satisfied by the observed *Q*_0_(*T*), …, *Q*_4_(*T*) and hence that at the level of cell populations averages the hypothesis of statistical independence between activations of distinct alleles *had to be rejected*.

### MM6. Maximum entropy model and dependency between alleles activations

Because full independence of the four alleles is not compatible with our experimental data, we computed, for each T, the joint probability *F*_*T*_ which minimizes dependency between alleles activation, and is still compatible with the observed *Q*_0_(*T*), …, *Q*_4_(*T*) via the linear relations imposed by Equation 1. For probability distributions verifying a set of linear constraints, a fairly generic principle is that minimizing dependencies is roughly equivalent to maximizing entropy. Recall that for any probability *F* = [*F*_0_, …, *F*_15_] on the set *S* = [*S*_0_, …, *S*_15_] of joint alleles activation, the entropy *Ent*(*F*) ≥ 0 of F is given by *Ent*(*F*) = − *F*_0_ *log*(*F*_0_) − *F*_1_ *log*(*F*_1_) − …. − *F*_15_ *log* (*F*_15_).

In Suppl. Materials 3, we apply this principle to compute the unique probability of joint alleles activation frequencies, denoted *F*_*T*_ = [*F*_*T*_(*S*_0_), …, *F*_*T*_(*S*_15_)] which has maximum entropy among all probabilities compatible with the observed frequencies *Q*_0_(*T*), …, *Q*_4_(*T*) via the five linear relations of Equation 1. Our formulas show that the maximum entropy joint probability F_**T**_ must have full symmetry, meaning that permutations of the alleles AL_1_, AL_2_, AL_3_, AL_4_ do not change the frequencies of their joint activation states. Indeed, we have the explicit formulas

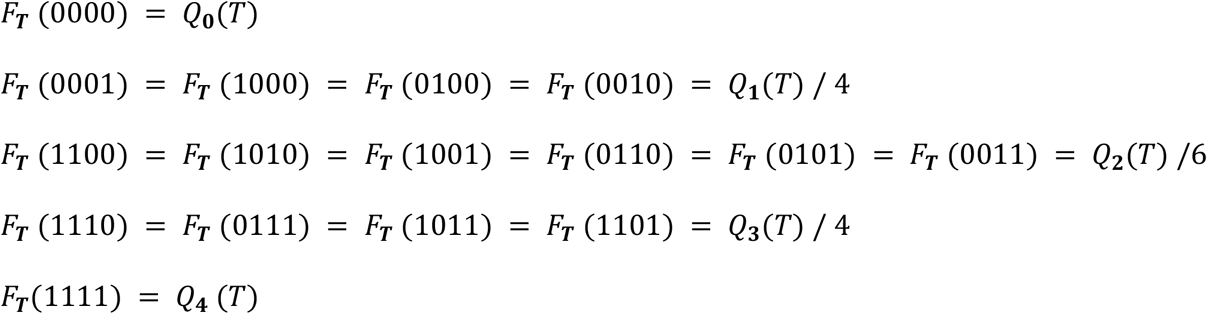

Since in our smFISH experimental images it is not possible to match specific alleles from cell to cell, full symmetry for the average probability *F*_*T*_ of joint alleles activations is a natural feature for compatibility with our image data.

Fix any single allele *AL*_1_. Due to the full symmetry of F_**T**_, the frequency

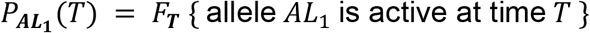

takes the same value for all four alleles, namely (see Suppl. Materials 3),

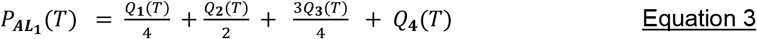

Consider now two alleles *AL*_1_ and *AL*_2_ in the same nucleus. At time T, the (random) state of *AL*_1_ is either “active” or “inactive”, which we denote by *AL*_**1**_ = 1 or *AL*_**1**_ = 0. Same remark for the random state of *AL*_**2**_. The mutual information 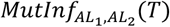 between the binary valued random states of *AL*_**1**_ and *AL*_**2**_ is given by the generic formula

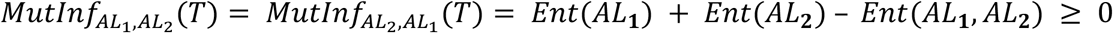

where “Ent” denotes the entropy of a probability distribution. Higher values of 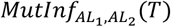 indicate higher dependency between the random activation states of *AL*_**1**_, *AL*_**2**_ under the probability *F*_*T*_. The maximum possible value of 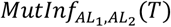 is 0.69, which can only be reached if there is a fully deterministic relation between the random activations of alleles *AL*_1_, *AL*_2_. But 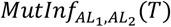 values higher than 0.02 already indicate some significant level of dependency between *AL*_1_, *AL*_2_. Conversely, values of 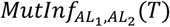 extremely close to 0 reflect near independence between the activation states of *AL*_1_ and *AL*_2_.

At time T, the pair (*AL*_1_, *AL*_2_) has 4 possible *joint states* (00), (01), (10), (11). Their frequencies *M*_*T*_00, *M*_*T*_01, *M*_*T*_10, *M*_*T*_11, are easily derived from the explicit expression (se equation 3) of the probability *F*_*T*_, which yields

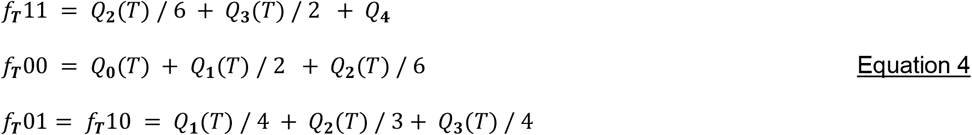

The probability 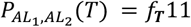 that *AL*_1_ and *AL*_2_ are simultaneously active at time T is hence given by

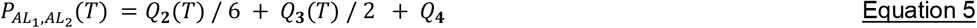

Then the joint entropy *Ent*(*AL*_**1**_, *AL*_**2**_) is given by

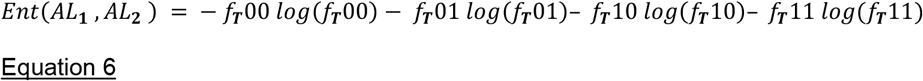

By definition of entropy, one has also

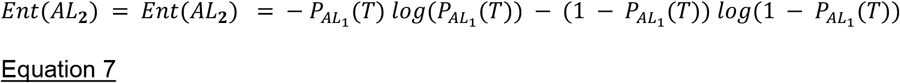

The *mutual information* 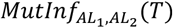 between the random activations of *AL*_**1**_ and *AL*_**2**_ can then be computed by

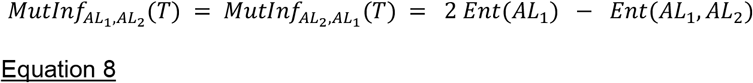

The equations 4,5,6,7,8 clearly express 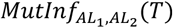 in terms of the 5 observed activations frequencies *Q*_0_(*T*), …, *Q*_4_(*T*). Due to the full symmetry of joint probability *F*_*T*_, the mutual information 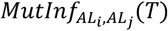 will be the same for all pairs of alleles *AL*_*i*_, *AL*_*j*_, and this common value *quantifies the average amount of activation dependency* between pairs of alleles at time T.

### MM7. Modeling the time course of GREB1 transcription frequencies observed at population level

Since genes transcriptions are strongly decorrelated from cell to cell, the random transcription bursts occurring among (for instance) the 400 cells of population *pop*(*T*) will be highly de-synchronized. Hence, averaging the random bursts that actual occur at fixed time T will clearly *smooth out* the impact of random bursts on the allele activation frequencies *Q*_***j***_(*T*) observed across *pop*(*T*). We have validated this point by simulations of 400 two-states stochastic models of GREB1 transcription in single cells, and averaging at each time T the nascent mRNA outputs of these independent 400 models. As expected, in population averaged transcription activity, short transcription bursts were essentially no longer identifiable. So, to emulate the time course of our population averaged GREB1 transcription data, we have introduced a population level stochastic model.

In this simplified model successive GREB1 transcriptions are initialized at random times *t*_**1**_ < *t*_**2**_ < … < *t*_***k***_. Each initialized transcription launches the elongation of a GREB1 mRNA molecule at fixed linear speed and is completed after a fixed *Mean Transcription Duration* MTD which for GREB1 is ≈ 44min. The random time intervals (*i*_***k***+**1**_ − *i*_***k***_) are assumed to be independent and to have the same *exponential density* with unknown *mean value A*. The successive occurrences *i*_***k***_ of transcription initializations define then a Poisson stochastic process. The complete nascent mRNA “*nas*_***k***_” generated by GREB1 transcription started at time *i*_***k***_ becomes fully visible at time (*t*_***k***_ + *MTD*). Both exonic and intronic parts of *nas*_***k***_, denoted *exonas*_***k***_. and *intnas*_***k***_, are naturally assumed to have exponential decay. The mean half-life of *exonas*_***k***_ is about 3 hrs, and hence does not impact the visibility of *nas*_***k***_ during the analyzed time-course (90min of E2 treatment). However, the mean lifetime of intronic *intnas*_***k***_ is known to be shorter than ≈ 35 min, and hence directly affects the first visibility time of *nas*_***k***_. The random lifetime of *intnas*_***k***_ from completion of *nas*_***k***_ to nearly full decay of *intnas*_***k***_ is assumed to have an *exponential density* with unknown fixed *mean value L*. Our population level stochastic model thus has only 3 key parameters {*A, L, MTD*}. To take into account the limits imposed by image resolution, we introduce another integer valued parameter, the unknown *Visibility Threshold* (*VTH*) such that nascent mRNA spots are detectable only if they contain at least *VTV* molecules (*VTH* = 2 was found after simulations).

Our *population level model* is easy to simulate, and we have fitted its 4 parameters {*A, L, MTD, VTH*} to each FV+E2 experiment by intensive simulations as outlined below. For E2 experiments with no FV pre-treatments, we must take account of actual GREB1 transcription activity occurring at low frequency in our cell populations during the last hour before the time *T* = 0 of E2 treatment. This requires the introduction of another parameter, namely the mean waiting time *A*^+^ between successive transcription initiations occurring *before* time *T* = 0.

### MM8. Simulations and model fitting

To fit our stochastic population level model to experimental data we performed intensive simulations to select the parameters {*A*^+^, *A, L, MTD, VTH*} providing the best quality of fit to our smFISH image data. These parameters were constrained to have naturally pre-defined ranges:

- mean transcription duration *MTD*: 40*min* < *MTD* < 50*min*, since RNA Polymerase II speed ≈ 2.5 kb/min and GREB1 length ≈110 kb
- mean lifetime *L* of nascent mRNAs: 5*min* < *L* < 35*min*, based on actinomycin D pulse-chase experiments
- mean waiting time A between rounds of transcription *after T* = 0 : 5*min* < *A* < 35*min*, based upon the on/off time ranges evaluated in (2)
- mean waiting time *A*^+^ < *A* between rounds of transcription *before T* = 0 : *A*^+^ < 60 *min*, based upon analysis of initial activations frequencies *Q*_**0**_(0), …, *Q*_**4**_(0). Note: *A*^+^ is used only for experiments with no FV pretreatment.

The minimum number *VTV* of molecules needed for reliable detection of nascent mRNA spots had to be crudely pre-calibrated by image analysis. As detailed in section MM9 below, and similarly to (2), we calculated the integrated intensities of mature, cytoplasmic mRNA spots to roughly evaluate the number of molecules per detected nascent mRNA spot. This yielded a *VTV* range between 1 and 10 molecules per spot.

These parameters ranges were discretized into finite grids with accuracies of 0.5 min to 1 min for all time variables. This gave us a multigrid of roughly 10^6^ possible parameter vectors *PAR* = {*A*^+^, *A, L, MTD, VTH*}. For each potential vector *PAR*, we performed a first set of 1000 simulations of our population level model. Each simulation outputs a random number *NAS* (*T*) of nascent mRNAs present at time T on a single virtual allele *AL*_**1**_. Among the 1000 simulated *NAS* (*T*), we compute the percentage 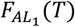 of integers *NAS* (*T*) which are larger than the visualization threshold *VTH*. Then 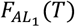 is the model generated frequency of detectable single allele activations. For each experiment and each vector *PAR* we then evaluate the quality of fit between model and data by the distance

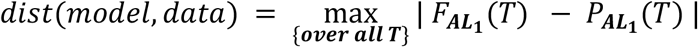

For the three FV+E2 experiments, the early values 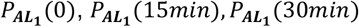 were practically 0 up to errors of estimations (3%), and the simulated 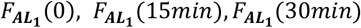 were identically zero since *MTM* was known to be of the order of 44 min. Therefore, for FV+E2 experiments, the distance between model and data was actually replaced by

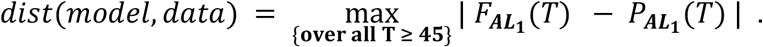

For each experiment, the best choices for the model parameters vector PAR are obtained by optimizing the quality of fit, i.e. by minimizing the distance *dist*(*model, data*).

We implemented the stochastic simulations of our population level model by the following algorithm. We first generate the random times *t*_***k***_ by standard simulation of the sequence of independent random waiting times (*t*_***k***+**1**_ − *t*_***k***_) having the same exponential density with mean *A*. Elongation of the nascent mRNA *nas*_***k***_ begins at *i*_***k***_ and is completed at time (*i*_***k***_ + *MTM*). The random lifetime *N*_***k***_ of *nas*_***k***_ is provided by a separately simulated sequence of independent random lifetimes *N*_***k***_ having the same exponential density with mean *L*. Then at each time T, the number of *nas*_***k***_ present at time T is determined by the number of *nas*_***k***_ such that *t*_***k***_ + *MTD* < *T* < *t*_***k***_ + *MTD* + *U*_***k***_. This simulation algorithm is naturally faster than the Gillespie algorithm used for more complex stochastic models.

Our Python simulation code is accessible in the publicly available software package on GitHub (https://github.com/smahmoodghasemi/BCM). This “brute force” approach to model fitting required intensive computing and was implemented on the “Sabine” multicore computing center at University of Houston. Once the simulations have been completed for 10^6^ models, this large set of simulations outputs can be re-used as a fixed massive lookup table for all our past or future experiments. After these simulations were successfully completed, we also outlined a more efficient computing approach which could be used in similar explorations for other genes. Namely, one can implement a multi-scale exploration starting with a cruder grid of parameters vectors, and then focus on finer mesh grids tightly centered around promising first level estimates.

### MM9. Estimation of number of molecules within detected nascent mRNA spots

For each *nascent* mRNA spot “*nas*” detected within the nuclei present in an image J, we compute the integrated exonic intensity *EXO*(*nas*) as the sum of image intensities *EXO*(*x*) over all exonic pixels x of “*nas*.” For each *mature* mRNA spot “*mat*” detected within the cytoplasm of all cells present in image J, we also compute the integrated exonic intensity *EXO*(*mat*) as the sum of image intensities *EXO*(*x*) over all pixels x of “*mat*.” We then compute the median *MED* of these integrated intensities *EXO*(*mat*) over all mature RNA spots detected in image J.

In the spirit of an approach explored in (2), the number *MOL*(*nas*)) of RNA molecules present within any *detected* nascent mRNA spot “*nas*” is crudely estimated by the 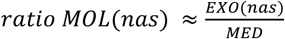 can only be a rough calibration of *MOL*(*nas*) since the observed *EXO*(*mat*) values have a high. This dispersion around their median *MED*, even at fixed time T. Nevertheless, to evaluate reasonable ranges for our model parameter *VTH* (Visualization Threshold) on any given image J, we have computed the low quantiles for the histogram of all *MOL*(*nas*) values extracted by computer analysis of image J. These low quantiles identified a potential range of 2 to 5 for the minimum number *VTH* of RNA molecules necessary to detect a nascent mRNA spot in our images. Since the error margins for these estimates were likely to be high, we simply assigned a much wider potential range of 1 to 10 for the unknown parameter *VTH*. After fitting of our population level to experimental data, our results reported above showed that the best estimate of *VTH* was always *VTH* = 2.

### MM10. 1. Estimation errors for frequencies Q_j_(T) and probabilities 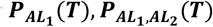

Tables given above and Table 6 in Supplemental Materials display the computed estimation errors for 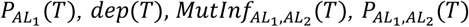. Here we outline how these errors are computed. Let *N*(*T*) = # cells in population *pop*(*T*). Use shorter notations 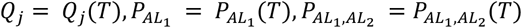. Let 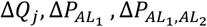 be the corresponding random errors of estimation. The 5’5 covariance matrix *COV*_***Q***_ of the 5 errors [Δ*Q*_0_, Δ*Q*_1_, …, Δ*Q*_4_] is classically given by

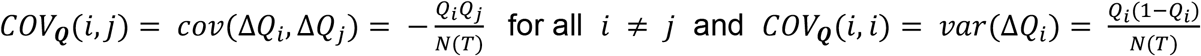

Denote *Q* the column vector [*Q*_0_; *Q*_1_; *Q*_2_; *Q*_3_; *Q*_4_], and for any matrix *H*, denote *H*^***tr***^ the transpose of matrix *V*. Due to Equations 3 and 5, the formulas for 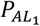 and 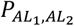 can be rewritten in matrix form as

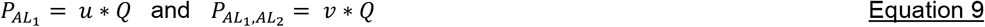

where *u* = [0, 1/4, 1/2, 3/4, 1] and *v* = [0, 0, 1/6, 1/2, 1]

Known statistical formulas then give the variances of 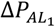 and 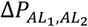 as

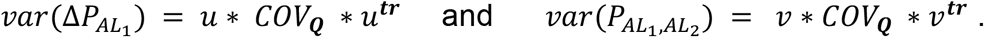

### MM10.2. Estimation errors for the dependency ratio 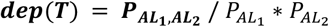

Due to Equation 9 we have 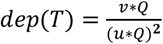 which is a nonlinear function *K*(*Q*) of *Q*. The variance of the random estimation error Δ*dep*(*T*) on *dep*(*T*) is then classically given by *var* [Δ*dep*(*T*)] = *w* * *COV*_***Q***_ * *w*^***tr***^ where w is the gradient *grad*_*Q*_ *K* of *K*(*Q*) with respect to *Q*. We have *w* = *grad*_***Q***_ *K* = (1 / *M* * *Q*)^**2**^ * *v* − 2 * [*v* * *Q* / (*u* * *Q*)^**3**^) * *u* which completes the computation of *var* [Δ*dep*(*T*)].

### MM10.3. Estimation errors for the mutual information 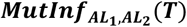

At any given time T, the pair of alleles (*AL*_1_, *AL*_2_) can be in one of their four joint activation states {11, 00, 01, 10} with corresponding joint probabilities *f*11, *f*00, *f*01, *f*10 given above by Equation 4. These formulas can be rewritten in matrix form as

*f*11 = *V*11 * *Q, f*11 = *V*11 * *Q, f*10 = *V*10 * *Q, f*01 = *V*01 * *Q*,

where the line vectors *Vij* are given by

*V*11 = [0, 0, 1/6, 1/2, 1] ; *V*00 = [1, 1/2, 1/6, 0, 0] ; *V*10 = *V*01 = [0, 1/4, 1/3, 1/4, 0]

The entropies *Ent* = *Ent*(*AL*_1_) = *Ent*(*AL*_2_) and *Ent*_12_ = *Ent*(*AL*_1_, *AL*_2_) are given by

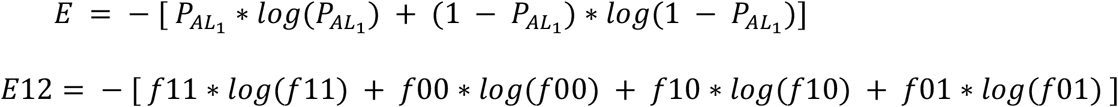

which can be rewritten as

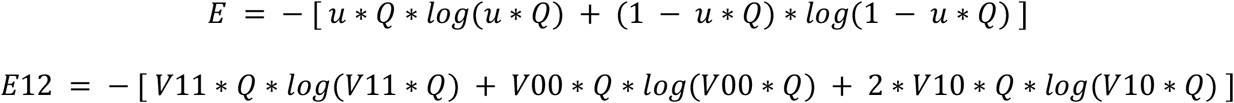

The information 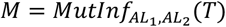, is then given by *M* = 2 * *Ent* − *Ent*12 which is a non linear function *M* = *G*(*Q*). The variance *vnasrob*(Δ*M*) of the random estimation error ΔM on M involves as above the gradient *grad*_***Q***_*G* of the function G(M) and is given by the formula

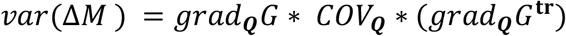

The preceding formulas directly yield

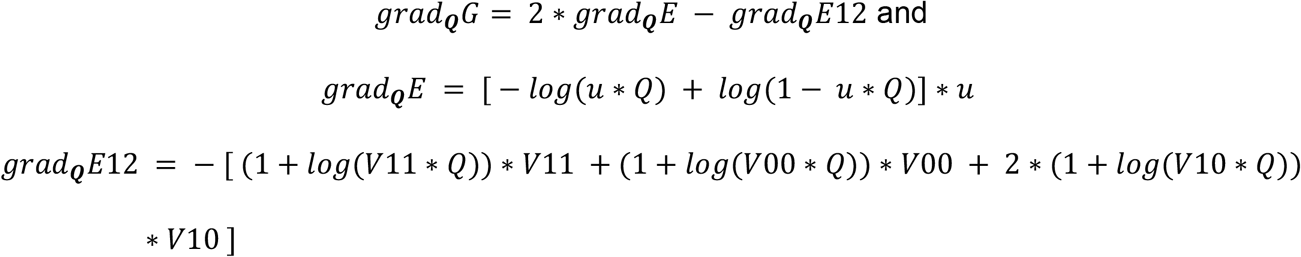

which clearly completes the computation of *vnasrob*(Δ*M*).

## ACKNOWLEDGMENTS

Imaging for this project was supported by the Integrated Microscopy Core at Baylor College of Medicine and the Center for Advanced Microscopy and Image Informatics (CAMII) with funding from NIH (DK56338, CA125123, ES030285), and CPRIT (RP150578, RP170719), the Dan L. Duncan Comprehensive Cancer Center, and the John S. Dunn Gulf Coast Consortium for Chemical Genomics. Modeling and computing research for this project at University of Houston were supported by CAMII and Baylor College of Medicine with funding from CPRIT (RP170719).

Intensive Computer Simulations for this project were implemented at the SABINE Cluster of RCDC (Research Computing Data Core, University of Houston) under a CPU-GPU computing time allocation to the University of Houston Department of Mathematics.

## Supplementary Materials: Algorithmics and Tables

### Suppl.Material 0: Tables for observed frequencies Q_0_(T) Q_1_(T) Q_2_(T) Q_3_(T) Q_4_(T)

For each one of our 6 experiments, image analysis at time T yields five key frequencies *Q*_*j*_(*T*). In cell population *pop*(*T*), *Q*_*j*_(*T*) is the frequency of nuclei exhibiting exactly “*j*” detected GREB1 nascent mRNA spots.

For two E2 experiments and two FV+E2 experiments, we display here the observed values of the *Q*_*j*_(*T*), listed as *percentages*. Similar results hold for our two other experiments.

Since *pop*(*T*) has size *N*(*T*) ≥ 400 cells, the estimation error on each *Q* (*T*) is less than 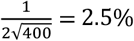.

E2 experiment #1
| T in minutes | $T = 0$ | $T = 15$ | $T = 30$ | $T = 45$ | $T = 60$ | $T = 75$ | $T = 90$ |
| --- | --- | --- | --- | --- | --- | --- | --- |
| $Q_0(T)$ | 77.0 | 67.3 | 53.5 | 39.6 | 30.0 | 26.0 | 22.0 |
| $Q_1(T)$ | 15.0 | 17.8 | 19.8 | 19.8 | 18.0 | 17.0 | 16.0 |
| $Q_2(T)$ | 5.0 | 9.9 | 15.8 | 19.8 | 21.0 | 19.0 | 18.0 |
| $Q_3(T)$ | 2.0 | 4.0 | 7.9 | 13.9 | 20.0 | 22.0 | 24.0 |
| $Q_4(T)$ | 1.0 | 1.0 | 3.0 | 6.9 | 11.0 | 16.0 | 20.0 |

E2 experiment #2
| T in minutes | $T = 0$ | $T = 15$ | $T = 30$ | $T = 45$ | $T = 60$ | $T = 75$ | $T = 90$ |
| --- | --- | --- | --- | --- | --- | --- | --- |
| $Q_0(T)$ | 45.5 | 37.6 | 31.0 | 26.0 | 19.8 | 14.0 | 11.9 |
| $Q_1(T)$ | 25.7 | 23.8 | 20.0 | 15.0 | 13.9 | 13.0 | 9.9 |
| $Q_2(T)$ | 14.9 | 18.8 | 20.0 | 19.0 | 19.8 | 20.0 | 18.8 |
| $Q_3(T)$ | 9.9 | 13.9 | 18.0 | 21.0 | 23.8 | 29.0 | 30.7 |
| $Q_4(T)$ | 4.0 | 5.9 | 11.0 | 19.0 | 22.8 | 24.0 | 28.7 |

FV+E2 experiment #4
| T in minutes | $T = 0$ | $T = 15$ | $T = 30$ | $T = 45$ | $T = 60$ | $T = 75$ | $T = 90$ |
| --- | --- | --- | --- | --- | --- | --- | --- |
| $Q_0(T)$ | 91.0 | 85.9 | 78.0 | 59.0 | 32.3 | 18.0 | 13.9 |
| $Q_1(T)$ | 6.0 | 10.1 | 13.0 | 15.0 | 15.2 | 13.0 | 9.9 |
| $Q_2(T)$ | 3.0 | 4.0 | 6.0 | 10.0 | 16.2 | 18.0 | 16.8 |
| $Q_3(T)$ | 0.0 | 0.0 | 2.0 | 10.0 | 20.2 | 27.0 | 29.7 |
| $Q_4(T)$ | 0.0 | 0.0 | 1.0 | 6.0 | 16.2 | 24.0 | 29.7 |

FV+E2 experiment #5
| T in minutes | $T = 0$ | $T = 15$ | $T = 30$ | $T = 45$ | $T = 60$ | $T = 75$ | $T = 90$ |
| --- | --- | --- | --- | --- | --- | --- | --- |
| $Q_0(T)$ | 90.1 | 88.0 | 84.7 | 71.7 | 41.6 | 23.0 | 23.8 |
| $Q_1(T)$ | 7.9 | 10.0 | 12.2 | 12.1 | 13.9 | 14.0 | 11.9 |
| $Q_2(T)$ | 2.0 | 2.0 | 2.0 | 6.1 | 14.9 | 19.0 | 15.8 |
| $Q_3(T)$ | 0.0 | 0.0 | 1.0 | 6.1 | 15.8 | 22.0 | 23.8 |
| $Q_4(T)$ | 0.0 | 0.0 | 0.0 | 4.0 | 13.9 | 22.0 | 24.8 |

### Suppl.Materials 1: Linear constraints on the average probability of joint alleles activations

For any fixed nucleus NUC_n_ of the population pop(*T*), each allele *AL*_*j*_, *j* = 1,2,3,4 can either be active (state “1”) or not (state “0”). There are 16 joint *alleles states* for *AL*_1_, *AL*_2_, *AL*_3_, *AL*_4_ naturally indexed by the first 16 binary numbers as follows

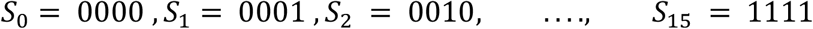

Denote *q*_***k***,***n***_ the probability that *NUC*_***n***_ exhibits exactly *k* activation spots at time T, and let *prob*_***n***_. be the joint probability of alleles activations in *NUC*_***n***_; we then have the basic probabilistic relations

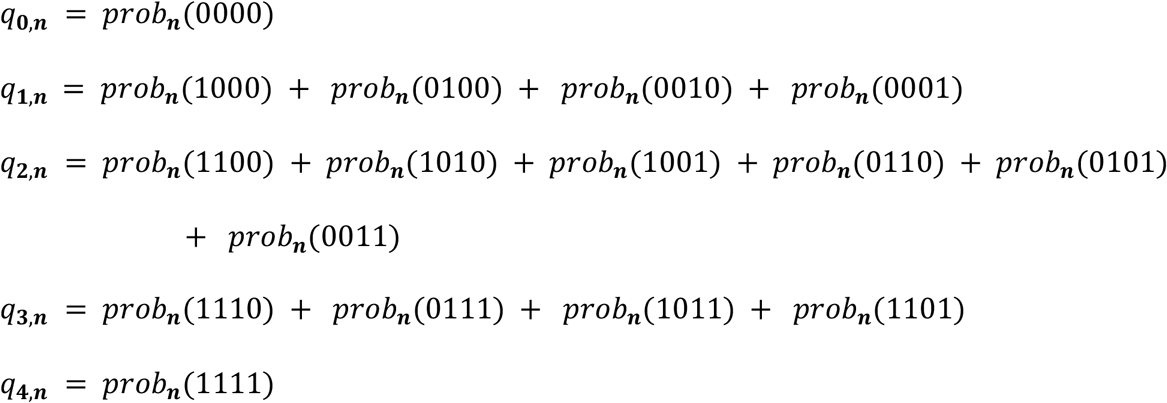

Note that *Q*_*k*_(*T*) is the average of the *q*_***k***,***n***_ over all nuclei *NUC*_***n***_ in *pop*(*T*), and that *F*_*T*_ is also the average of the *prob*_***n***_ over *i*. After averaging over *i*, the preceding linear relations yield

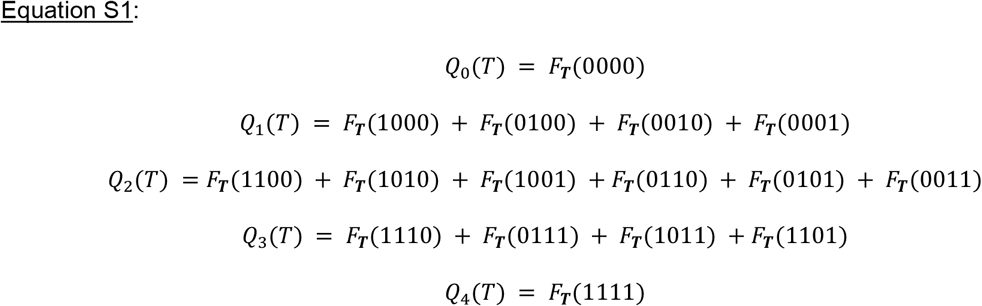

### Suppl.Materials 2: Impact of independence on probabilities of joint alleles activations

Fix the time T. Under the average probability *F*_*T*_ of joint alleles activations just defined, let *M*_*j*_ be the probability that allele *AL*_*j*_ is activated at time T. Then *h*_*j*_ = 1 − *M*_*j*_ is the frequency of non-activation for *AL*_*j*_. Assume temporarily that under the joint probability *F*_*T*_, the random activations of 0 *robrob* 1, independence of alleles activations implies

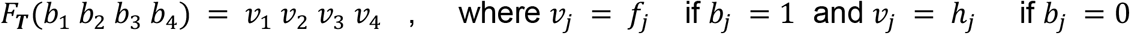

Combining these product formulas with the linear relations of equation S1 proves that the nuclei activation frequencies *Q*_*k*_ = *Q*_*k*_(*T*) must verify the five formulas

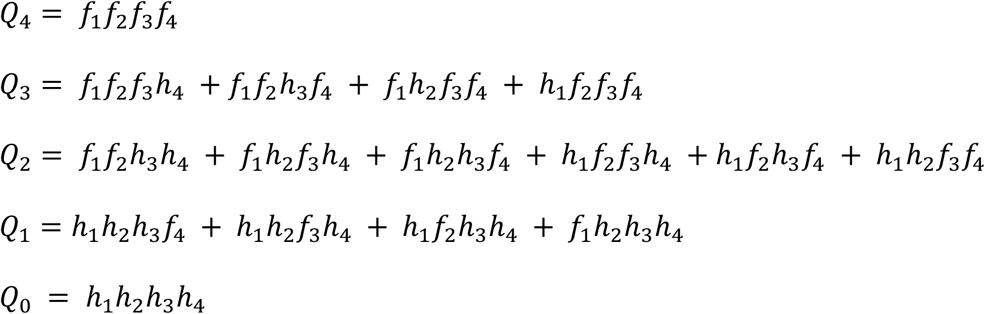

Divide the first four of these equations by the last one and set *z*_*j*_ = *M*_*j*_ / *h*_*j*_ for *j* = 1, 2, 3, 4, to obtain

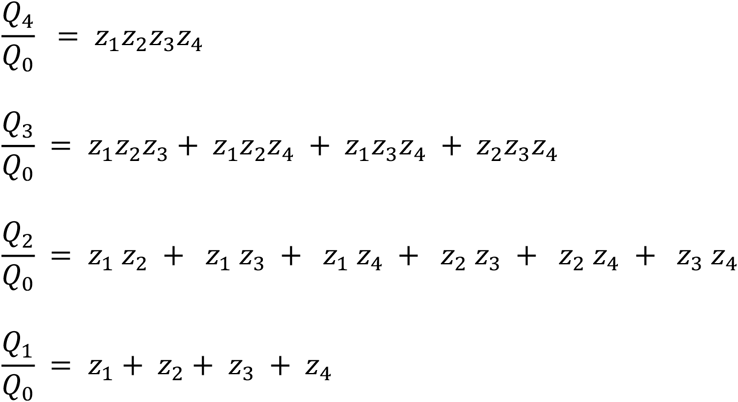

These formulas classically imply that *z*_1_, *z*_2_, *z*_3_, *z*_4_ must be the four roots of the polynomial equation

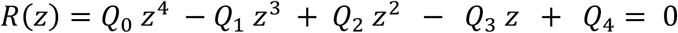

Hence independence of allele activations under the joint probability *F*_*T*_ forces the polynomial *PAR*(*z*) to have 4 *positive real valued solutions z*_1_*z*_2_*z*_3_*z*_4_.

The relations 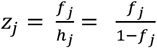 then imply that the unknown probabilities *f*_2_, *f* _2_, *f* _3_, *f*_4_ are given by

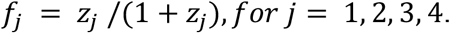

The independence of alleles activations thus requires the polynomial *PAR*(*z*) to have all its roots real and positive, a condition which imposes *very restrictive polynomial constraints* on the 5 observed frequencies *Q*_*j*_(*T*) for each time T. In particular, each one of the 10 pairs *Q*_*i*_(*T*), *Q*_*j*_(*T*) with *i* < *j* must verify very specific polynomial inequalities. For instance, the pairs *Q*_3_, *Q*_4_ and *Q*_1_, *Q*_0_ must verify

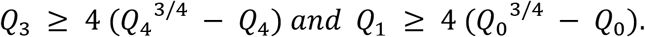

In all our experiments and at all positive times T, the polynomial *R*(*z*) with coefficients *Q*_0_(*T*), …, *Q*_4_(*T*) derived for image analysis of smFISH data actually did NOT have four positive and real valued roots. This led us to *reject the alleles independence hypothesis* for the population average probability *F*_*T*_ of joint alleles activations

### Suppl.Material 3: Maximum Entropy under Constraints

Fix time T and denote F for short the probability *F*_*T*_ = [*F*_0_ *F*_1_ … *F*_15_] on the finite state space *S* = [*S*_0_, …, *S*_15_] of joint alleles activations. The entropy *Ent*(*F*) is given by

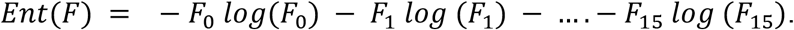

The five frequencies *Q*_*j*_ = *Q*_*j*_(*T*) are known and fixed. We know that F must verify the 5 linear relations given by equation S1, which can be rewritten with more compact notations as

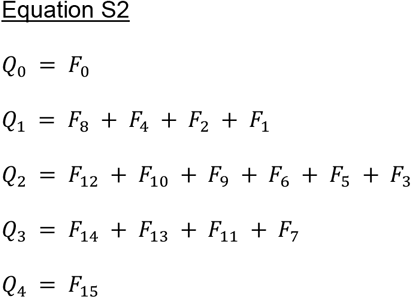

To seek a probability *F* maximizing *Ent*(*F*) under the 5 linear constraints of equation 2, and the linear relation {*F*_0_ + … + *F*_15_ = 1}, we introduce 6 Lagrange multipliers *L*_1_… *L*_6_. The partial derivative *M*_*min*_ of *Ent*(*F*) with respect to *F*_*min*_ is equal to [−1 − *log*(*F*_*min*_)]. The 16 classical Lagrange conditions for optimization under constraints are then

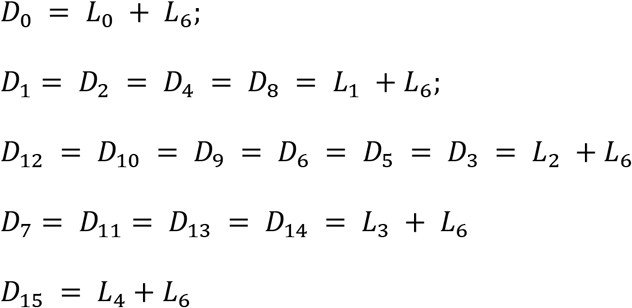

Since *M*_*min*_ = −1 − *log*(*F*_*min*_) the 5 preceding equations show that

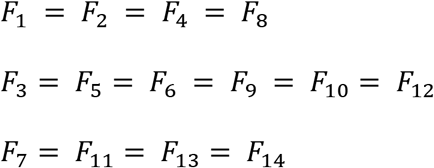

Reporting these equalities in Equation S2 yields directly

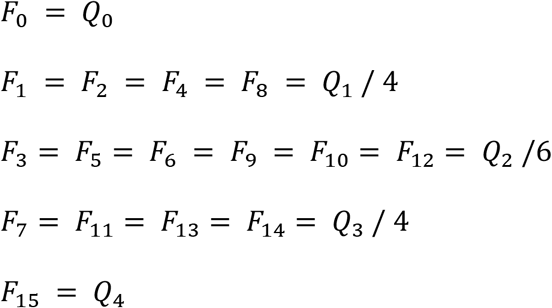

This fully determines *F* = *F*_*T*_, and also proves that *F*_*T*_ has *maximum symmetry, i*.*e*., is *unchanged by any permutation* of the alleles *AL*_1_, *AL*_2_, *AL*_3_, *AL*_4_. Indeed, the preceding expressions obtained for the probability *F* = *F*_*T*_ can be rewritten

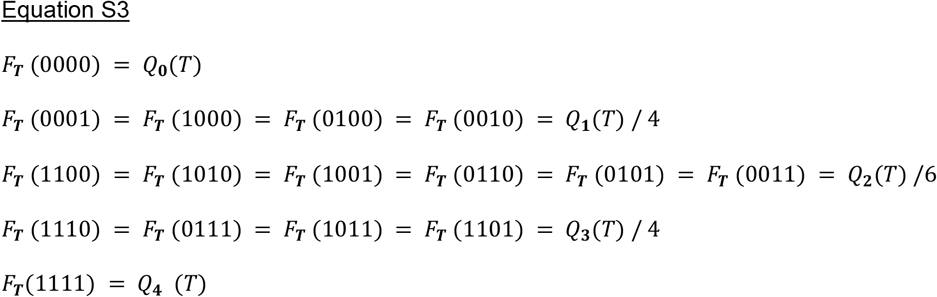

Due to the maximum symmetry of *F*_*T*_, the probability 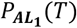 that allele *AL*_1_ is active at time T has the same value for all 4 alleles. By definition 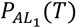 is given by the sum

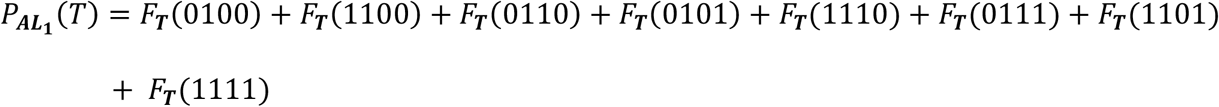

The explicit formulas S3 just obtained for F_T_ yield then

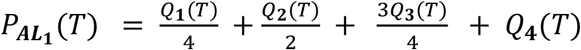

A similar computation provides the joint probability 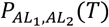 as was outlined in “Methods”.

